# Patient-Derived Air-Liquid Interface Forebrain Organoids Reveal Functional Synaptic Deficits in Schizophrenia

**DOI:** 10.64898/2026.06.01.729183

**Authors:** Lucrezia Criscuolo, Pia Jensen, Helle Bogetofte, Sissel I. Schmidt, Fadumo A. Mohamed, Lene A. Jakobsen, Marie S. Øhlenschlæger, Henriette R. S. Frederiksen, Fangyuan Li, Elif Bayram, Michael E. Benros, Jonthan Brewer, Barbara L. Lind, Phillip J. Robinson, Kristine Freude, Martin R. Larsen

## Abstract

Schizophrenia (SCZ) is a severe and debilitating neurodevelopmental disorder with lifelong impact on everyday life. Disruptions in synapse functions play a key role in its complex and poorly understood etiological and pathological mechanisms.

Here, we investigated both the molecular composition and the spontaneous and stimulated functional properties of synapses in neural organoids from SCZ individuals. Air-liquid interface forebrain organoids (ALI-FOs) were generated from induced pluripotent stem cells (iPSCs) derived from three individuals with SCZ and three healthy controls. At day 170 synaptosomes were enriched and analyzed by data-independent acquisition mass spectrometry to profile the proteome, alongside with TMT-labeled phosphoproteomics both before and after acute KCl-induced depolarization. In parallel, we characterized the PTMome of the surrounding cellular environment, comprising phosphorylation, peptides with free and reversibly modified cysteines, and sialylated N-linked glycopeptides. Functional glutamatergic and GABAergic activity was assessed using calcium imaging to capture spontaneous neuronal signaling.

Both conditions exhibited mature synaptic structures, while growth cones were observed only in SCZ-derived ALI-FOs, indicative of ongoing or delayed synaptogenesis. Proteomic analysis of synaptosome preparations revealed 358 differentially regulated proteins between SCZ and controls and 125 phophoproteins with altered phosphorylation, which clustered into three major categories: (1) synaptogenesis and synapse signaling; (2) cytoskeleton and cell junctions; (3) growth cone dynamics and neurite outgrowth. Analysis of the PTMs in the surrounding cellular environment revealed regulation of key regulatory mechanisms in 526 proteins, supporting the synaptic alterations observed. Notably, components of the Wnt signaling pathway were consistently dysregulated across both the synaptosome preparation and the PTMome in SCZ-derived ALIFOs as compared to controls. Depolarization-induced phospho-signaling revealed SCZ-specific response enriched in synaptic vesicle trafficking pathways.

Together, these findings provide new insights into early synaptic alterations in SCZ, highlighting changes not only in protein composition, but more in protein regulatory mechanisms underlying synaptic signaling.

## Introduction

Schizophrenia (SCZ) is a severe and complex neuropsychiatric disorder affecting more than 20 million people worldwide^1,2^. It is characterized by a combination of various debilitating symptoms, comprising behavioral and cognitive dysfunctions (e.g., hallucination, delusion, psychosis, disorganized speech) and negative symptoms (e.g., lack of emotional expression or response, anhedonia, social withdrawal), which chronically impair everyday life^2–5^.

Typically, SCZ manifests in late adolescence or early adulthood, but it is widely accepted that the pathological processes originate during the early brain development, in the so-called prodromal period^6–9^. Understanding the underlying molecular mechanisms for disease development in the early stages is essential for elucidating SCZ pathogenesis, which remains poorly understood and challenging to investigate due to crucial experimental limitations. Access to human brain tissue, especially from the early developmental phases, is highly limited by technical and ethical constraints^9–11^. As a result, longitudinal studies are rare, and no cellular models exist that can effectively recapitulate the temporal gap between clinical symptom onset and initial disease mechanisms.

Although SCZ has a high heritability (over 60%), no single causal gene has been identified^2,3,12^. Instead, the disorder has been associated with a heterogeneous genetic background, involving several risk genes and more than 200 loci over the past years, suggesting a multifactorial nature of SCZ, where genetic and environmental factors interplay^4,13–15^. This heterogeneity is probably reflected in the wide range spectrum of symptoms, pathological mechanisms, and treatment responses, posing a continuous challenge to our understanding of the molecular basis of SCZ^3,16^.

The lack of representative samples and models further complicates the SCZ research field^14^. Recently, neural organoids (NOs) have emerged as valuable *in vitro* models to study the early stages of brain development^17^. With their three-dimensional cytoarchitecture, self-organization, self-development and diverse cell composition, NOs resemble key features of brain structure and biology, overcoming several limitations of previous models^11,18,19^. The advancing of protocols enables the generation of region-specific NOs (e.g., forebrain, midbrain, hindbrain), as well as more sophisticated *in vitro* systems including functional neuronal networks, multi-brain region integration, and transplantation in host animals^18–22^. Additionally, NOs can be generated directly from patient-derived induced pluripotent stem cells (iPSCs), maintaining the individual genetic background and enabling the modeling of complex polygenic neurodevelopmental disorders, such as SCZ and autism spectrum disorder (ASD)^17,23^. NOs have already been used to study physiological and pathological brain development^11,17^, model neurodegenerative diseases (e.g., Parkinson’s disease, Alzheimer’s disease)^24–26^, test pharmacological agents (e.g., valproic acid)^27^, and investigate the effects of infectious and exogenous factors (e.g., Zika virus, SARS-CoV-2)^20,28^. SCZ has been successfully modeled in NOs, revealing novel risk genes (e.g., PODXL, PTN), cell-specific neuropathology, disrupted neurogenesis and altered electrophysiological activity^9,11^. Despite its potential, NO-based research is in its early stages, and organoids remain experimental models that still require standardized protocols, interpretations, and validation frameworks.

In the past years, multiple hypotheses have been proposed to explain SCZ pathogenesis, including the dopamine, glutamate, and GABA hypotheses^14,29^. More recently, converging evidence has highlighted the central role of synaptic disturbance in SCZ^12,30^. Genomic and proteomic studies have associated several genes with synaptic pathways involved in synapse formation, organization, and function (e.g., SNAP91, SHANKs, DLGAP2, GRIN2A and GRIA3)^12,14,15,31^. Postmortem brain analyses have consistently reported numerous structural abnormalities, among which reduced spine density and neuronal size^14,32–35^. Neuroimaging studies further supported synaptic disturbance in SCZ brains, revealing impaired neural circuitry, disrupted connectivity, and abnormal synchrony^36,37^.

It is hypothesized that synaptic involvement in SCZ may arise from either impaired synaptogenesis, during early brain development, or excessive synaptic pruning coinciding with clinical symptom onset^5,8,35,38^. However, experimental and clinical evidence of synapse disturbance in SCZ remains incomplete and needs further investigations.

In this study, we investigated synaptic structures and their proteome in early brain development in SCZ using our newly developed approach for the enrichment and characterization of synapse structures from ALI-FOs^39^. We generated air-liquid interface forebrain organoids (ALI-FOs) from three individuals with SCZ and three healthy controls, culturing them for 170 days to promote synaptic maturation. Synaptic structure fractions were then enriched from these organoids and subjected to in-depth, unbiased proteomic and phosphoproteomic profiling to uncover SCZ-associated synaptic alterations. To further assess synaptic function, we evaluated the phospho-signaling responses of ALI-FO-derived synaptic structures following acute KCl-induced depolarization and analyzed the spontaneous glutamatergic and GABAergic calcium activity within the ALI-FO system. In parallel, we conducted a comprehensive analysis of multiple post-translational modifications (PTMs) - including phosphorylation, peptides with free cysteines, peptides with reversibly modified cysteines, and sialylated N-linked glycopeptides - in the surrounding cellular environment to capture additional layers of functional alterations in SCZ ALI-FOs, supporting the synaptic alterations.

Notably, while both conditions exhibited mature synaptic structures, growth cones were observed only in SCZ-derived ALI-FOs at day 170. Proteomic and phosphoproteomic analysis of synaptic structures, and PTMome characterization of the surrounding environment revealed consistent alterations across three interconnected functional categories: synaptic formation and signaling; cytoskeleton rearrangements and cell junctions; and growth cone dynamics and neurite outgrowth. Remarkably, several components of the Wnt signaling pathway were depleted in SCZ-derived synaptic structures, and consistently dysregulated within the PTMome. Despite the limited sample size, these findings offer new insights into the molecular and functional synaptic disturbance in the early brain development in SCZ.

## Methods

### Participant recruitment

Individuals with SCZ and healthy controls were recruited according to the ethical approval (j.no: S-20220037 from the Danish Health Research Ethics Committees in the 33 Southern Region of Denmark) and the data protection approval (j.no: P-2022-765 from The Danish Data Protection Agency) from psychiatric clinics in the Capital Region of Denmark and via the Danish website, Forsøgsperson.dk, respectively. The inclusion criteria for recruitment were as follows: participants must be between 18 and 50 years of age and must come from the same community. All individuals went through psychiatric evaluations including WHO Schedules for Clinical Assessment in Neuropsychiatry (SCAN, version 2.1), covering the last 4 weeks to confirm the schizophrenia diagnosis (according to the International Classification of Diseases, Tenth Revision (ICD10) F20) and to exclude any psychiatric disorders for healthy controls.

### Sample collection, PBMC isolation and episomal reprogramming

Venous blood samples were collected at the research facilities of the Copenhagen Research Centre for Biological and Precision Psychiatry, Mental Health Centre Copenhagen (Copenhagen University Hospital) and immediately transported to the Department for Veterinary and Animal Science (University of Copenhagen) for peripheral blood mononuclear cell (PBMC) isolation. A 1:1 dilution of 34 mL of blood sample and Dulbecco′s Phosphate Buffered Saline (PBS-/-; Sigma Aldrich) supplemented with 2% Fetal bovine serum (FBS; Fisher Scientific) was performed and then slowly added to 15 ml Lymphoprep™ (Stem Cell Technologies) contained into 2 separate SepMate™-50 columns (Stem Cell Technologies). After a centrifugation at 1,200xg for 10 min at room temperature, the top layer of the sample was transferred to a 50 mL falcon tube, diluted with PBS supplemented with 2% FBS, and centrifuged again. This step was performed twice to ensure a pure fraction of PBMCs.

Then, PBMCs from each participant were used to establish hiPSCs using episomal reprogramming. 4x 10⁶ PBMCs were cultured in PBMC media consisting of StemPro-34 SFM (Fisher Scientific) media supplemented with 100 ng/mL Human Stem Cell Factor Synthetic Peptide (Peprotech), 100 ng/mL Recombinant Human Flt3-Ligand (Peprotech), 20 ng/mL Interleukin-3 (Cell Guidance system), 1 ng/mL Recombinant Human GM-CSF (Peprotech), 2 mM L-glutamine (Thermo Fisher Scientific) and 1% penicillin/streptomycin (P/S) (Sigma Aldrich) in low-attachment 6-well plates for 2 days. Then, 1 million cells were counted and cultured in PBMC media without P/S for 1 day and reprogrammed into hiPSC using a modified protocol from Konala V.B.R. et al^40^. Thus, PBMCs were transfected with 1 µg/µl of pCXLE-hOCT3/4-shp53 (Addgene) pCXLE-hSK (Addgene), pCXLE-hUL (Addgene) pCXB-EBNA1 (Addgene) in a ratio 1:1:1:1 using the P3 Primary Cell 4D X Nucleofector Kit L and the Nucleofector Lonza Machine (nucleofection program EO-100; BioNordika). Then, after 3 min at RT, 300 µL of PBMC medium with 1% P/S and 10 µM ROCK inhibitor (ROCKi) (Sigma Aldrich) were dropwise added to the cells and incubated for 10 min at 37°C with 5% CO_2_. Then, the cells were cultured on growth factor reduced Matrigel (Corning) in PBMC medium, and the medium was carefully changed with fresh PBMC media with 1% P/S the day after. After 3 days of reprogramming, 50% of the medium was changed to stem cell media composed of mTeSR1 medium (Stem Cell Technologies) with 0.1% P/S and 250 µM Sodium Butyrate (Sigma Aldrich). The stem cell medium was then changed every second day until colonies emerged. Approximately 16 days after reprogramming, colonies were manually picked and cultured on Matrigel with stem cell media and supplemented with 10 µM ROCK inhibitor. Ultimately, a minimum of 6 colonies per cell line were expanded to generate hiPSCs clones and banked. The hiPSC clones were assessed by the pluripotency markers nanog, OCT-4 and TRA-1-60 through immunocytochemistry (not shown here).

### ALI-FOs generation

The iPSC lines were maintained in mTeSR Plus medium (Stem Cell Technologies) on growth factor reduced Matrigel and were used to generate ALI-FOs according to the STEMdiff Dorsal Forebrain Organoid Differentiation Kit (Stem Cell Technologies) and to the protocol from Giandomenico S.L. et al.^19^ with the following modifications. On day 0, after dissociation of iPSCs into single cells, 3×10^6^ cells were seeded per well in AggreWell 800 24-well plate (Stem Cell Technologies) in 2 mL FBO Formation medium supplemented with 50 μM ROCK inhibitor (Sigma-Aldrich). On day 6, approximately 60 embryoid bodies (EBs) per cell line were transferred to low-adherent 6-well plates with 2 mL/well FBO Expansion Medium and placed on an orbital shaker at 57 rpm (INFORS HT Celltron) until day 40-45 to avoid organoid fusions. Between days 40-45, ALI-FOs were generated by slicing around 25-30 organoids per cell line into 300 µm ALI-FO sections using a vibratome (VT1000S, Leica). Then, 5-6 ALI-FOs were placed per Millicell® cell culture inserts (Millipore) with Serum-Free Slice Culture medium beneath (SFSCM), consisting of Neurobasal medium (Thermo Fisher Scientific), supplemented with 1:50 (v/v) B-27 (Thermo Fisher Scientific), 1x (v/v) GlutaMAX (Thermo Fisher Scientific), 1:10 (v/v) P/S (Thermo Fisher Scientific) and 1:500 (v/v) Fungin (InvivoGen). Additionally, from day 48 ALI-FOs were freshly supplemented with 0.2 µM L-ascorbic acid (Merck), 20 ng/mL BDNF (Stem Cell Technologies) and 20 ng/mL GDNF (Stem Cell Technologies). After two weeks, the cell culture medium was gradually exchanged from SFCM to BrainPhys Neuronal medium (containing SM1 Supplement and N2 Supplement-A; Stem Cell Technologies) over a 3-week period, using the following scheme: one week of 75% SFCM and 25% BrainPhys mixture, followed by a 50%:50% mixture in the second week, and finally a 25% SFCM and 75% BrainPhys mixture during the third week.

All cells and organoid cultures were maintained with 5% CO_2_ at 37°C.

### Synaptic structure enrichment

Synaptosomes were isolated following the protocol described by Ohlenschlaeger, M.S., et al. ^39^ with minor changes. For each cell line, ALI-FOs at day 170 were collected, pooled and ice-cold sucrose/EDTA buffer (0.32 M sucrose, 1 mM EDTA, 5 mM Tris base, pH 7.4) was added in a ratio of 750 mL/100 mg of ALI-FOs. The samples were then homogenized with 6 up-and-down strokes at 700 rpm using a glass tissue grinder (15-ml Potter-Elvehjem type Teflon-glass tissue grinder; Wheaton). Subsequently, the homogenate was centrifuged at 800xg for 15 min at 4°C to pellet nuclei and cellular debris (P1 fraction), while the supernatant (S1 fraction) was collected. The P1 fraction was subsequently resuspended in ice-cold sucrose/EDTA buffer, and the centrifugation repeated a second time, so the new S1 fraction was combined with the previous one. After measuring the protein concentration using a nanophotometer N60 (Implen), each S1 fraction was divided into 2 replicates of 1.5 mL each, corresponding to approximately 3 mg of proteins. Finally, the synaptosomes were isolated using a centrifugation at 5,000xg for 10 min at 4°C, resulting in the collection of P5000 fractions.

### Synaptosome stimulation

P5000 synaptosome fractions were resuspended in 100 µL of 37°C-warm HBK control buffer (118 mM NaCl, 4.7 mM KCl, 20 mM HEPES, 1.18 mM MgSO_4_, 1.2 mM CaCl_2_, 0.1 mM Na_2_HPO_4_, 10 mM glucose, 25 mM NaHCO_3_, 10 mM Pyruvate, pH 7.4) and incubated at 37°C for 1 h. One sample of P5000 fraction per cell line was then stimulated for 15 sec using 100 µL of high KCl HBK buffer (147.7 mM KCl, 20 mM HEPES, 1.18 mM MgSO_4_, 1.2 mM CaCl_2_, 0.1 mM Na_2_HPO_4_, 10 mM glucose, pH 7.4), while the other replicate went through the same procedure but with HBK control buffer. The reaction was stopped by snap-freezing in liquid nitrogen and the samples stored at-70°C until further use.

### Sample preparation for TMT labelled analysis

Sodium Deoxycholate (SDC) was added to the KCl-stimulated and non-stimulated P5000 fractions to a total of 2%. The samples were sonicated using a probe sonicator (Q125 Sonicator Ultrasonic Homogenizer, Homogenizers, Atkinson, NH, US) for 2×20 sec at 40% amplitude. After sonication, the samples were ultracentrifuged at 100,000g for 1 h. After centrifugation, the supernatant was transferred to low binding Eppendorf tubes. The pellets were resolubilized by probe sonication in 100 mM HEPES, pH 8.5 containing 10% SDC for 2×20 sec at 20% amplitude and subsequently diluted to 5% SDC. The soluble and pellet protein concentration was then measured using the nanophotometer N60 (Implen). A total of 100µg of protein were taken out from the soluble fractions and 50 µg from the pellet fractions for subsequent TMT labeling. Proteins were reduced by adding 10 mM dithiothreitol (DTT; Sigma) for 30 min followed by alkylation with 20 mM iodoacetamide (IAA, Sigma) for 30 min in the dark at RT. After alkylation, the IAA reaction was stopped by the addition of 5 mM DTT. Protein digestion was performed by incubating the samples with (0.04 units per mg of proteins) Lys-C (Lysyl endoproteinase; Wako Chemicals) for 1 h at 37°C, followed by an overnight incubation with 1:20 (w/w; enzyme/substrate ratio) trypsin (modified trypsin, in-house synthesized) at 37°C. The day after, each sample was labelled with 0.25 mg of TMTpro™ (Tandem Mass Tags; Thermo Fisher Scientific) for 1.5 h at RT according to manufacturer instructions, resulting a two 12plex TMT samples. Then the samples from the same fraction were combined in equal ratios and the TMT reaction was quenched with 5% hydroxylamine (ThermoFisher) for 15 min. The SDC was subsequently precipitated from the combined samples using 2% (v/v) formic acid (FA, Merck) and centrifuging at 20,000xg for 15 min.

### Enrichment of phosphopeptides

Phosphopeptides from P5000 fractions were enriched as described previously^41^. Briefly, peptides were dissolved in loading buffer (80% acetonitrile (ACN; VWR), 5% Trifluoroacetic acid (TFA; Merck), 1 M glycolic acid (Sigma)) and incubated shaker at the highest shaking speed (approx. 2,000 rpm) for 15 min at RT with titanium dioxide (TiO_2_) beads (GL Sciences Inc) in a beads-to-peptides ratio of 1 mg per 100 µg of peptides. A second incubation of 10 min with half of the previous amount of TiO_2_ beads was performed, and the supernatant, containing non-modified (NM) peptides, was collected and dried by vacuum centrifugation. Phosphopeptides were eluted from the TiO_2_ beads using 5% ammonia water, pH 11 (Merck) and incubated overnight with PNGase F (New England BioLabs) and sialidase A (Prozyme) for deglycosylation. The two fractions (phosphopeptides/deglycopeptides and non-modified peptides) were purified and desalted by in-house-made reversed-phase microcolumns made of Oligo R3 Resin (Applied Biosystems) and Empore SPE disks C18 (Sigma) packed into a p200 pipette tips. Desalted peptides were eluted using 60% ACN, 0.1% TFA and dried by vacuum centrifugation prior to high pH reversed phased fractionation.

### High-pH reversed-phased fractionation

Non-modified, phosphopeptide, deglycopeptide, and RmCys samples were resuspended in 20 mM ammonium formate (pH 9.3), centrifuge at 14,000xg for 10 min at RT, and transferred into a microtiter plate. Then, the samples were fractioned using a Dionex Ultimate 3000 HPLC system (Thermo Scientific) and an Acquity UPLC®-Class CSHTM C18 column (Waters). For phosphopeptide, deglycopeptide, and RmCys fractions a total of 12 concatenated high pH fractions were collected using the following gradient of solvent B (80% acetonitrile, 20% of 20 mM ammonium formate): 2-50% in 74 min, 50-70% in 10 min, and 70-95% in 5 min, hold at 95% for 10 min, return to 2% over 17 min. The non-modified fractions were fractionated in 20 concatenated fractions using the following gradient of solvent B: 2-38% in 115 min, 38-95%in 15 min, hold at 95% for 10 min, return to 2% over 1 min.

All fractions were lyophilized and resuspended in 0.1% FA for LC-MS/MS analysis.

### Sample preparation for label free analysis

A small aliquot (200-400 ng) of the trypsin/Lys-C digests derived from P5000, P1 and S1 fractions was taken for DIA analysis.

### Sample preparation for PTM enrichment

The P1 pellets were resuspended in 0.1% SDS, sonicated and boiled at 95°C for 10 min. Then, the samples were incubated for 1 h with 5 mM CysPAT to block the free cysteines^42^, and subsequently ultracentrifuged at 100,000xg for 1 h at RT to separate SDS soluble from insoluble material (membranes including potential remaining active zones). The insoluble fraction was resolubilized in 100 mM HEPES, pH 8.5 containing 10% SDC, probe sonicated for 2×20 sec at 40% amplitude and diluted to 5% SDC. The soluble fraction was subjected to 10 KDa spin filters where the SDS was replaced with 3% SDC, and 100 mM HEPES, pH 8.5. Protein concentrations were measured using the N60 instrument.

The proteome and PTMome (defined here as phosphorylation, free cysteines, reversibly modified cysteines and sialylated N-deglycopeptides) were analyzed essentially as described in the MOSAIC-PTM protocol^43^.

### Liquid chromatography-mass spectrometry in data dependent acquisition (DDA)

All high pH RP fractions were dissolved in buffer A (0.1% FA) and analyzed by nano LC-ESI-MS/MS using an EASY-nLC (Thermo Fisher Scientific) with buffer A (0.1% FA) and buffer B (95% ACN, 0.1% FA) connected online to an Orbitrap Exploris^TM^ 480 mass spectrometer (Thermo Fisher Scientific, US) (Phosphopeptides and depalmitoylated peptides) or an Orbitrap Eclipse Tribrid mass spectrometer (Thermo Fisher Scientific). The samples were loaded onto an in-house made pulled emitter analytical column (100 μm inner diameter packed with Reprosil-Pur 120 C18-AQ, 1.9 μm beads (Dr. Maisch GmbH)). The phosphopeptide/deglycopeptide fractions and the non-modified peptide fractions were eluted using increasing buffer B (95% ACN, 0.1% FA) from 2 to 25% in 100 min and then from 25 to 40% B for 20 min.

For the fractions containing the non-modified and modified peptides a full MS spectrum was obtained in the mass range of 350–1200/1500, in the Orbitrap with a resolution of 120,000 full-width half maximum (FWHM), a maximum injection time of 50 ms, and an Automatic Gain Control (AGC) target value of 10^6^. Hereafter, the peptides were automatically selected for MS/MS for a 3 sec cycle time using higher energy collision dissociation (HCD) with normalized collision energy (NCE) setting at 34, resolution of 45,000 FWHM, AGC target value of 10^5^ ions and maximum injection time of 150 ms (phosphopeptides/deglycopeptides).

### Liquid chromatography-mass spectrometry in data independent acquisition (DIA)

All label-free samples were analyzed by nano LC-MS/MS, using a Vanquish NEO system (Thermo Fisher Scientific) coupled with Orbitrap Astral mass spectrometer (Thermo Fisher Scientific). The samples were loaded onto a two-column system with a 5 mm x 300 µm Acclaim™ PepMap™ 100 C18 HPLC trap column (5 µm, Thermo Scientific) and an in-house packed 20 cm x 100 µm ID Reprosil-Pur C18-AQ RP analytic column (1.9 µm; Dr. Maisch GmbH, Germany).

The peptides were separated with an increasing gradient of solvent B (95% ACN/0.1% FA) at a flow rate of 300 nL/min. The gradient was as follows; 2-40% in 20 minutes, 40-100% in 1 minute, then 100% for 5 minutes. Full MS1 scans were acquired with an automatic gain control (AGC) target of 500%, scan range at 400-1000 and an Orbitrap resolution of 240,000. For the DIA analysis an isolation window of 2 Da was used with a total of 299 windows over the mass range. A HCD normalized collision energy of 32 was used with a filling time of 5ms and an MS cycle time of 0.6 sec. Normalized AGC target value of 500% was used for all DIA scans.

### Transmission Electron Microscopy (TEM) imaging

P5000 fractions were fixed in 3% glutaraldehyde in 0.1 M sodium phosphate buffer (78 mM Na_2_HPO_4_, 22 mM NaH_2_PO_4_, pH 7.2) for 1 h at 4°C. The fixing solution was then replaced with 0.1 M sodium phosphate buffer and samples were stored at 4°C.

For fixation, the buffer was removed, and fractions were embedded in 2% agar solution. Solidified agar was cut into 2×2 mm blocks that were prepared for dehydration by washing two times in 0.1M Na-phosphate buffer for 5 min, incubated in 1% Osmium/0.1 M Na-phosphate buffer for 1 h followed by two washing in MilliQ water for 5 min. Samples were dehydrated by gradually transferring the samples from ethanol to 1.2 propylenoxid (PO) to Epon using the following order: 50% EtOH – 10 min, 70% EtOH – 10 min, 96% - 10 min, 99% EtOH – 2×20 min, PO – 10min, PO/Epon 2:1 – 20 min, PO/Epon 1:1 – 20 min, PO/Epon 1:2 – 20 min, Pure Epon – overnight. Next day, the dehydrated samples were cast in molds with Epon and curated at 60°C for 24-48 h.

Curated samples were sectioned in 0.1µm sections and contrasted using 2% uranyl at 50°C for 30 min, washed 7 times in MilliQ water, incubated with lead citrate for 7 min at room temperature and washed 7 times in MilliQ water. Images were obtained at x8500, x17500 and x22000 magnification.

### Viral transduction

On day 155, 6 ALI-FOs per cell line were transduced for calcium imaging recording. Briefly, 3 ALI-FOs per cell line were individually incubated with a 1:1 ratio (1x 10^10^ viral genome in BrainPhys Neuronal medium) of pAAV.Syn.NES-jRGECO1a.WPRE.SV40 (a gift from Douglas Kim & GENIE Project; Addgene viral prep. 100854-AAV1) and pENN.AAV.CamKII.GCaMP6f.WPRE.SV40 (a gift from James M. Wilson; Addgene viral prep 100834-AAV1), while other 3 ALI-FOs per cell line were individually transduced with 1:1 ratio (1x 10^10^ viral genome in BrainPhys Neuronal medium) of pAAV.Syn.NES-jRGECO1a.WPRE.SV40 and AAV-mDlx-jGCaMP8f-WPRE (a gift from Loren Looger; Addgene viral prep. 176753-AAV1). The ALI-FOs were incubated for 24 h at 37°C with 5% CO_2_ and then the medium was changed to regular BrainPhys Neuronal medium. ALI-FOs were maintained at 37°C with 5% CO_2_ for 2 weeks, before proceeding with calcium imaging recording.

### Calcium imaging

All experiments were carried out on a laser scanning multiphoton excitation system (FluoView FVMPE-RS, Olympus) paired with a Mai Tai HP Ti:Sapphire laser (Millenia Pro, Spectra-Physics) using a 25x magnification and 1.05 numerical aperture water-immersion objective (Olympus). A dichroic mirror split emitted light with a 500-600 nm band-pass, then the transmitted light was sent to an optical filter with a 489-531 nm band-pass and ultimately to multi-alkali photomultiplier tube (PMT), corresponding to the GCaMP6f or jGCaMP8f fluorescent signal. At the same time, the light reflected through the dichroic mirror was sent to 601-657 nm band-pass filter, and then to a high-sensitivity gallium arsenide phosphide (GaAsP) PMT, corresponding to the jRGECO1a fluorescent signal. The two additional detectors, one GaAsP PMT and one multi-alkali PMT, were used to collect reference signals to correct for background noise. Images were collected as a single plane by bidirectional scanning using a resonant scanner, with a maximum frame size of 512 x 512 pixels and a frame bit depth of 10 bit. Images were sampled at a minimum of 40 Hz, from at least 50-60 nm depth from the tissue surface. For each sample, a minimum of 6 field of views (FOVs) were acquired, where individual images were recorded for approximately 2 min, for 3 consecutive times, with 1 min-break to prevent phototoxicity. Laser emission was set to a wavelength of 1000 nm and the objective was immersed in 37°C-warm BrainPhys Neuronal medium.

The images were then analyzed in ImageJ (version 2.14.0/1.54f).

### Protein identification and quantification – DIA and DDA data

The raw data from DIA-MS analyses were first converted into HTRMS file format using HTRMS Converter (Biognosys, version 19.9) and then processed using Spectronaut® 19 (Biognosys) in direct DIA analysis. The data were searched against the Uniprot human database (uniprot_sprot_2024-01-01_HUMAN), with the following parameters: as enzymes both trypsin and Lys-c; carbamidomethyl (C) as fixed modification and acetyl (protein N-term) and oxidation (M) as variable modifications; a maximum of 2 missed cleavages; *exclude single hit proteins* was activated. Statistical analysis was performed in Spectronaunt® 19, using unpaired t-test, 0.58 as log_2_ fold change cut off, and 0.05 as q-value threshold.

TMT raw data were processed in Proteome Discoverer (v. 2.5.0.400, Thermo Fisher Scientific), using SEQUEST HT searched against the Uniprot human database. The search was performed using the following settings: as enzymes Trypsin full; a maximum of 2 missed cleavages; precursor mass tolerance of 10 ppm and fragment mass tolerance of 0.05 Da; TMTpro (K), TMTpro (N-term) and Carbamidomethyl (C) was set to static modifications except for the P1 fractons. For the searches of the P5000 phosphorylation/deglycosylation data the dynamic modifications included were Phosphorylation (S, T, Y) and Deamidation (N). For fraction P1 the data were searched with variable modification Phosphorylation (S, T, Y), Deamidation (N), SIA (C) and Carbamidomethyl (C).

Peptide identifications were filtered against a 1% false discovery rate cut-off using the Percolator algorithm^44^. For the non-modified peptide dataset, peptide identifications were assembled into protein groups. Quantification was performed based on the intensities of the TMT reporter ions.

Only proteins with two or more unique peptides were considered for further analysis in the non-modified group. Additionally, a protein was considered quantified only if it was quantified in at least 50% of the samples within the same condition. While in glycopeptide annotation, peptides with 2 or more deamidations were filtered out, as well as peptides not presenting the NXST motif. In addition, potential glycoproteins were searched in Uniprot database in search of known glycosylations; if any was annotated, the protein was kept for further analysis.

Statistical testing was performed in Proteome Discover using built-in t-test, and 0.05 as adjusted p-value cut off, and 0.58 as log_2_ fold change cut-off. Modified peptides that resulted differentially regulated were manually checked for spontaneous deamidation, missed cleavages, ambiguous identification and PTM assignment.

Gene Ontology (GO) term enrichment analysis was performed in R (version 4.4.0; R Core Team, 2024) using clusterProfiler R package^45^, or SynGO database^46^. Data visualization was carried out using ggplot2^47^ in R, the Omics Visualizer app^48^ in Cytoscape (version 3.10.3)^49^ and Biorender (https://BioRender.com).

### Immunohistochemistry (IHC) labeling and imaging

On day 170, 3 ALI-FOs were collected and fixed in 4% paraformaldehyde (Thermo Scientific) in PBS (Gibco) for 1 h at room temperature. After the incubation, the samples were washed three times in PBS and then soaked in 30% sucrose in PBS at 4°C for at least one night, or until embedding. ALI-FOs were then incubated in 7.5% gelatin (Sigma-Aldrich), 10% sucrose (Sigma-Aldrich) in PBS for 15 min at 37°C, and then embedded in the same gelatin/sucrose solution. The blocks were first solidified at 4°C for 15-20 min, then snap-frozen in-30/-50°C ethanol and stored at-70°C until sectioning.

The embedded ALI-FOs were sectioned in 20 µm sections on a Leica CM1860 cryostat at - 20°C, placed on microscope slides (Thermo Fisher Scientific) and stored at-20°C.

Then, the slices were hydrated with washing buffer (0.1-0.2% saponin (Sigma-Aldrich) in 1xPBS), incubated in quenching buffer (∼ 2 g of ammonium chloride (Sigma-Aldrich) in 30 mL of PBS) for 10 min and then washed twice with washing buffer. Subsequently, the slices were incubated in blocking buffer (5% donkey (VWR) or goat serum (Life technologies) in washing buffer) for 1 h, and then with primary antibodies in blocking buffer overnight at 4°C. After the incubation, the samples were washed 3x 10 min with washing buffer and incubated with secondary antibodies in blocking buffer for 2 h at RT in the dark. The slices were then washed 3x 10 min with PBS and mounted with DAPI-containing ProLong Diamond Antifade mountant (Invitrogen). Samples were stored in the dark at 4°C until image acquisition.

Images were acquired using a Nikon AX confocal/multiphoton microscope operated with NIS 5.4. Quadband detection and analyzed using ImageJ (version 2.14.0/1.54f).

| **Primary Antibody** | **Company** | **Cat. No.** | **Dilution** |
| --- | --- | --- | --- |
| **Rabbit anti-β tubulin** | Cell signaling | 5568 | 1:2000 |
| **Rabbit-anti vGlut1** | Synaptic Systems | 135303 | 1:1000 |
| **Mouse anti-β tubulin** | Sigma | T8660 | 1:2000 |
| **Rabbit anti-SYN1** | Abcam | ab254349 | 1:400 |
| **Rabbit anti-GFAP** | Dako | Z0334 | 1:3000 |
| **Rabbit anti-TOMM20** | Abcam | ab186735 | 1:250 |

| **Secondary Antibody** | **Company** | **Cat. No.** | **Dilution** |
| --- | --- | --- | --- |
| **Alexa Fluor 647 goat-anti-rabbit** | Invitrogen | A21245 | 1:1000 |
| **Alexa Fluor 488 goat anti-rabbit** | Life technology | A11008 | 1:1000 |
| **Alexa Fluor 568 goat anti-mouse** | Life technology | A11004 | 1:1000 |
| **Alexa Fluor 488 donkey anti-mouse** | Life technology | A21202 | 1:1000 |
| **Alexa Fluor 647 donkey anti-rabbit** | Life technology | A31573 | 1:1000 |

## Results

### Enrichment of synaptic structures from ALI-FOs

To investigate the molecular alterations in synapses in SCZ pathogenesis, we generated ALI-FOs from iPSCs from three individuals with SCZ and three healthy controls. The organoids were cultured for 170 days to ensure the formation and maturation of functional synapses and neuronal networks. ALI-FOs from the two conditions exhibited no macroscopic or morphological differences during the development. Subsequently their cytoarchitecture was evaluated by the expression of neuronal, synaptic, astrocytic and mitochondrial markers - SYN1, VGLUT1, β-TUB, GFAP and TOMM20 (Figure S1).

At day 170, synaptic structures were enriched from a pool of ALI-FOs (approximately 130-160 per cell line) using a differential centrifugation protocol optimized for ALI-FOs^39^. The quality of the synaptic structure fractions was assessed by TEM, which enabled the evaluation of both structure integrity and stage of synaptic maturation. As shown in Figure 2D, mature synaptic structures were observed in both CTL- and SCZ-derived fractions, including clear synaptic vesicles (SVs), mitochondria, and specialized areas (e.g., postsynaptic density (PSD)). Notably, all SCZ-derived synaptic structure fractions also contained structures resembling growth cones - developing neurite structures, characterized by filopodium, a more homogeneous cytoplasm, an almost complete lack of SVs, and absence of specialized regions.

**Figure 1.**
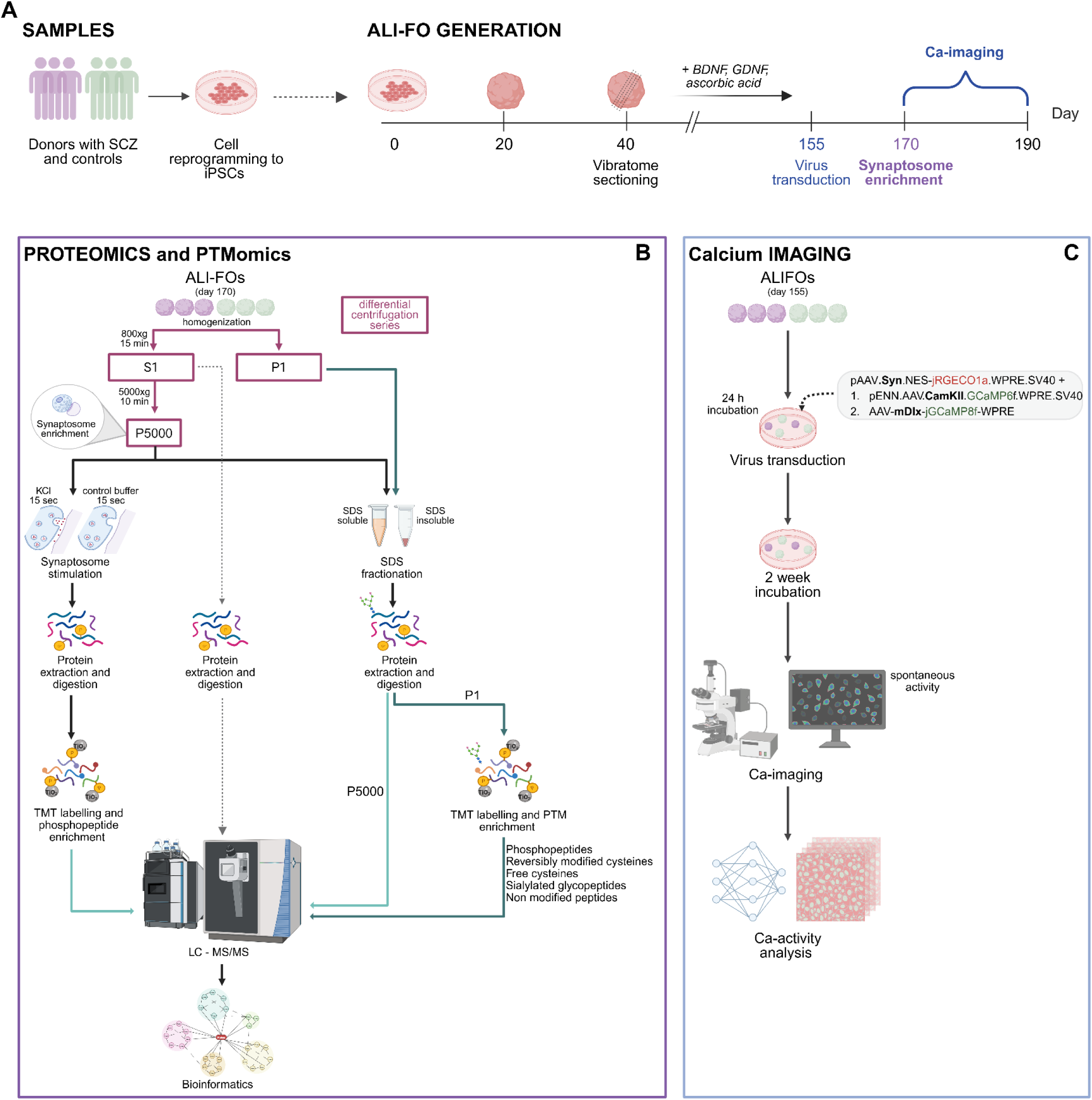
Schematic overview of the workflow. **A.** Briefly, the experimental setup starts with the generation of air-liquid interphase forebrain organoids (ALI-FOs) from induced pluripotent stem cells (iPSCs) derived from 3 individuals with schizophrenia (SCZ) and 3 healthy controls. **B.** At day 170, synaptosomes (P5000 fraction) are enriched from ALI-FOs using a differential centrifugation series and an aliquot was subjected to KCl stimulation. Mass spectrometry-based proteomics was performed on various fractions (P5000, P1, S1) from the differential centrifugation series. **C.** In parallel, calcium imaging was conducted on 6 ALI-FOs per cell line using two combinations of virus to distinguish glutamatergic and GABAergic neuronal activity.

**Figure 2.**
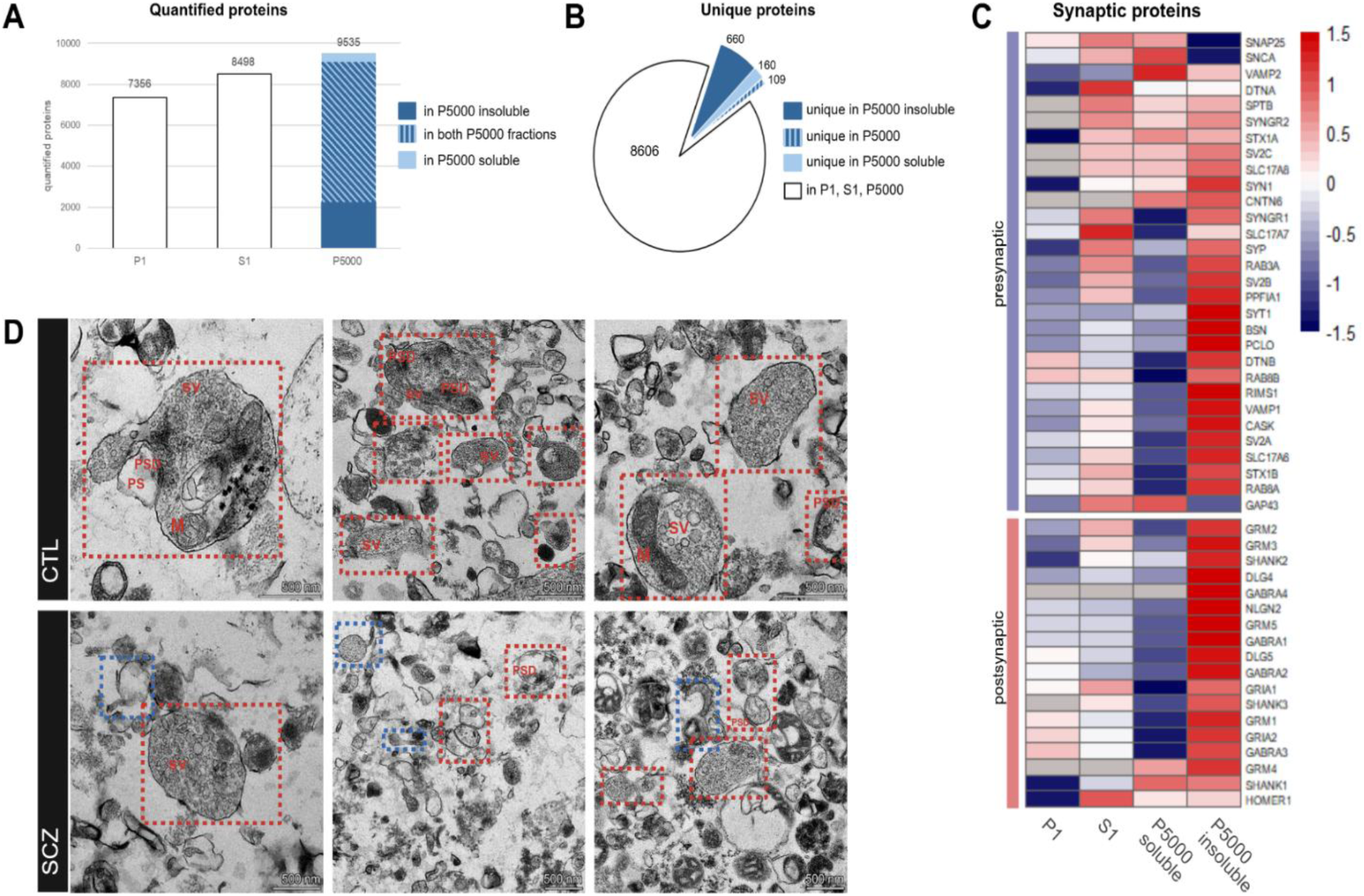
A. Bar plot showing the number of quantified proteins in the P1, S1 and P5000 fractions. **B.** Pie chart illustrating the proteins detected across all fractions (P1, S1, and P5000), and those unique to the P5000 fraction. **C.** Heatmap displaying the normalized abundance (Z-score) of synapse-specific proteins in the different fractions. **D.** TEM images of P5000 fractions derived from CTL and SCZ ALI-FOs, showing the presence of mature synaptosome (red). In SCZ images growth cones (blue) are also visible. Synaptosome components are labeled as follows: synaptic vesicles (SV); mitochondria (M); postsynaptic density (PSD); postsynapse (PS).

To characterize the molecular composition of synapses in SCZ and CTL, P5000 fractions containing synaptic structures were analyzed using large-scale data independent acquisition (DIA) mass spectrometry (MS), alongside with the initial crude fractions derived from the homogenate - P1 and S1 - for comparison. Given the architecture of synaptosomes - containing a detergent insoluble active zone (AZ) -, P5000 samples were further fractionated into SDS-soluble and SDS-insoluble fractions as described previously for enriched synaptosomes from mice^50^ to enhance proteomic coverage.

The use of SDS allows protein separation based on their chemical-physical properties, especially facilitating the detection of membrane-associated proteins – e.g., the ones in the AZ in synaptosomes.

Our MS-based analysis quantified 7356, 8498 and 9535 proteins in P1, S1 and P5000 fractions, respectively (Figure 2A). Within the P5000 samples, 9099 proteins were quantified in the SDS-insoluble fraction and 7286 in the SDS-soluble, with 6850 were shared between them. Notably, over 900 proteins were uniquely quantified in the synaptic structure-enriched fraction (Figure 2B), underscoring its relevance for profiling the synaptic protein composition. Additionally, to assess synaptic enrichment, the abundance of synapse-specific proteins was compared across P1, S1 and P5000 fractions. As shown in Figure 2C, the majority of both pre-and post-synaptic markers, including SV proteins (e.g., SV2B, SYP), active zone (AZ) components (PCLO, BSN), and neurotransmitter receptors (GABRA1-4, GRIA1), were enriched in at least one of the P5000 fractions.

Finally, across all investigated fractions, more than 98% of proteins were quantified in both SCZ-and CTL-derived ALI-FOs, revealing a near-complete overlap in protein identification between groups.

### Synaptic structure proteome

Less than 4% of quantified proteins were differentially regulated in SCZ-derived synaptic structures compared to controls (q-value < 0.05, log2FC <-0.58 or > 0.58), corresponding to a total of 358 proteins. Among these, 221 were identified in the SDS-insoluble fraction and 183 in the SDS-soluble fraction, with 46 in common between the two fractions. Of the total, 165 proteins were downregulated and 193 upregulated in SCZ.

To gain functional insight into these proteomic alterations observed in SCZ, we first annotated the differentially regulated proteins using SynGO, a manually curated synapse-specific database^46^. This analysis revealed that 50 proteins mapped to unique SynGO terms (Figure 3A), highlighting a strong involvement of synaptic structures in SCZ. These annotations included key presynaptic components, such as CADPS2, CNTN5, SYT12, as well as postsynaptic elements (e.g., DLG2, NEURL1, NRP1). Notably, several SV proteins associated with SV structure and cycle (e.g., SV2B, SLC17A7, PLD1, SYT12) were significantly regulated, in addition to regulators of neurotransmitter receptor level and localization (e.g., LPAR1, SST, HPCA, TMEM108), suggesting a potential disturbance of synaptic signal in SCZ. Several proteins implicated in growth cones and growth cone guidance - such as SEMA3E, involved in axon guidance signaling, NRP1, mediating semaphorin activity, and CNTN5, associated with neurite outgrowth – were also differentially regulated in SCZ.

**Figure 3.**
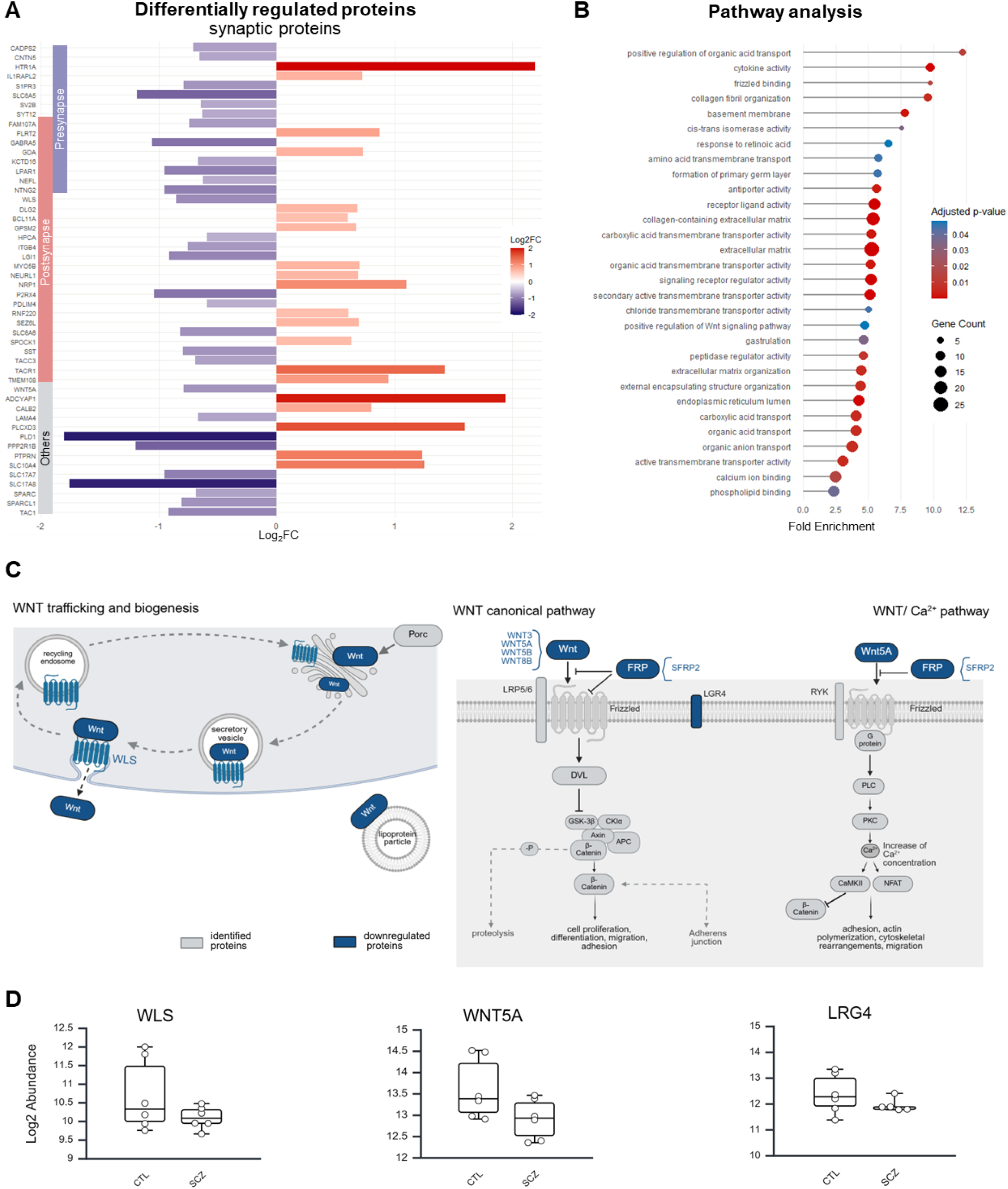
A.Plot showing synapse-specific proteins that are differentially regulated in SCZ synaptosomes compared to CTL. **B.** Pathway analysis of differentially regulated proteins in SCZ synaptosome fractions. **C.** Schematic overview of the WNT signaling pathway, highlighting differentially regulated proteins in SCZ synaptosome fractions. **D.** Boxplots of selected WNT pathway proteins showing differential expression between SCZ and CTL synaptosome fractions.

Finally, pathway analysis of the significantly regulated proteins, with a particular focus on the downregulated proteins in SCZ (Figure 3B), revealed enrichment in calcium ion binding, including SCIN, ANXA2, ANXA4, SPARC, HPCA, and S100B, and the Wnt signaling pathway. Specifically, Wnt pathway components depleted in SCZ included key regulators of Wnt trafficking and biogenesis, such as WLS, as well as ligands and modulators of both canonical and non-canonical pathways (WNT5A, WNT5B, WNT3, WNT8B, SFRP2, and LGR4), suggesting a potential alteration of Wnt signaling at the SCZ synapses (Figure 3C-D).

### The synaptic structure phosphoproteome

Given the fundamental role of PTMs in regulating protein biological activity, localization, and interactions^51,52^, the phosphoproteome of SCZ-and CTL-derived synaptic structures was investigated. To enhance quantification reliability through the analyzed conditions, the proteins extracted and digested from the synaptic structures were first labeled with TMT allowing multiplexing, and subsequently the phosphopeptides were enriched using TiO_2_ chromatography.

From a total of 10864 quantified phosphopeptides, corresponding to 2997 proteins, 125 were differentially regulated in SCZ. Of these, 53 were upregulated and 72 downregulated. Interestingly, as illustrated in Figure 4A, most of the dysregulated phosphoproteins fall into three main, and often interconnected, functional categories: (1) synapse organization and signaling; (2) cytoskeleton and cell junctions; (3) growth cones and axonogenesis (Supplementary Figure S2). Several key mediators of neuronal development and connectivity establishment were significantly affected. For instance, neuromodulin (GAP43) - a major marker of growth cones - plays a fundamental role in axonal and dendritic filopodium formation. Similarly, DCX (neuronal migration protein doublecortin), is crucial in promoting neural migration and interaction, as well as structural remodeling. DCC (netrin receptor DCC), mediates attractive guidance signaling in axonal growth cones. Additionally, other notable phosphoproteins included MARCKS, which modulates cytoskeletal dynamics and contributes to axon development, neurite initiation and outgrowth; SYN1 (synapsin I), a regulator of axonal outgrowth and synaptogenesis; and DTNA (dystrobrevin alpha) implicated in synapse formation and stabilization.

**Figure 4.**
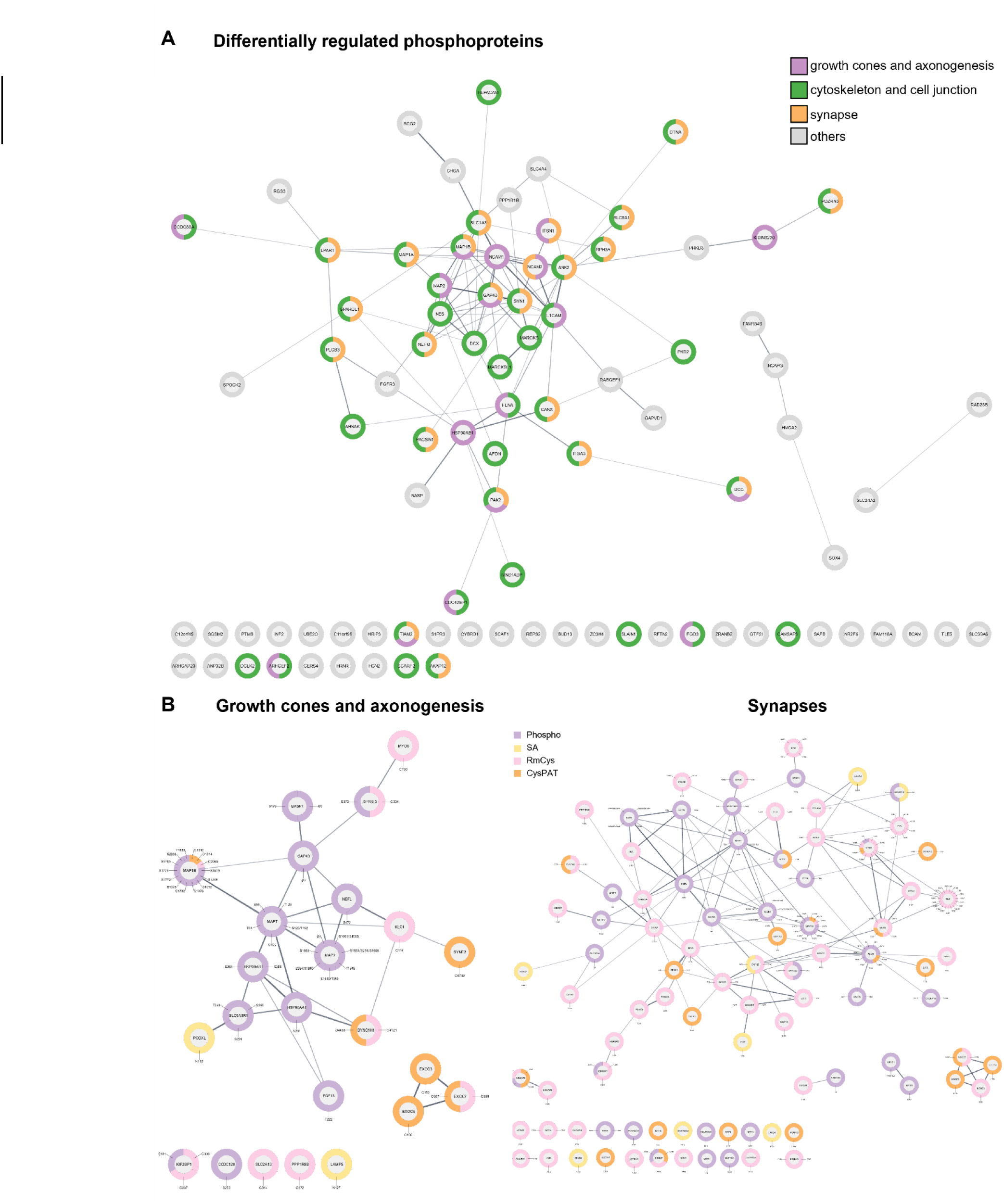
A.String network of differentially regulated phosphoproteins in SCZ synaptosomes compared to CTL. Proteins associated with growth cones and axonogensis (violet), cytoskeleton and cell junction (green), and synapses (orange) are highlighted. **B.** String networks of growth cones and neurite outgrowth-related proteins with differentially regulated post-translationally modified peptides detected in the P1 fraction. Modifications include sialylation (SA, yellow), phosphorylation (Phospho, violet), reversed modified cysteine (RmCys, pink), and free cysteine (CysPAT, orange)

### PTMome of the P1 fraction

Building upon the proteomic and, particularly, phosphoproteomic alterations observed in SCZ-derived synaptic structures, we extended our investigation to the P1 fraction - the initial pellet obtained from the differential centrifugation protocol. This fraction includes both the surrounding cellular environment, encompassing a broader cellular content from ALI-FOs not captured in the synaptic structure fractions, and additional synaptic structures. It is not surprising that certain synaptic populations - e.g. smaller or structurally distinct synaptic terminals along the neurites - are not efficiently released upon homogenization in hypotonic buffer and co-isolated through the differential centrifugation protocol, especially not from ALI-FOs. Analysis of this fraction could enable the identification of complementary molecular changes in synapses in SCZ. To enhance general proteomic coverage and increase the coverage of detergent-insoluble proteins, the P1 fraction was subjected to further separation into SDS-soluble and SDS-insoluble fractions, as performed for P5000. Based on previous studies^9,11^, major differences in overall protein abundance between SCZ and CTL are not expected. Instead, we hypothesized that alterations in biological activity of proteins, mediated by PTMs, might be more informative. Therefore, using our recently developed PTMomics workflow^43^, we identified and quantified several PTMs from P1 fraction, including phosphorylation, peptides with free cysteines (CysPAT), peptides with reversibly modified cysteines (RmCys), and formerly sialylated (SA) N-linked glycopeptides, in addition to non-modified proteins. In total, we quantified 4973 phosphopeptides, 1720 formerly SA N-glycopeptides, 7734 CysPAT peptides, 14025 RmCys peptides - corresponding to 1997, 818, 3174, and 5421 proteins, respectively - alongside 5799 non-modified proteins (Supplementary Figure S3). Statistical analysis identified significant changes in SCZ samples in 245 phosphopeptides (71 in SDS-insoluble and 176 in SDS-soluble), 44 formerly SA N-glycopeptides (19 in SDS-insoluble and 30 in SDS-soluble), 121 peptides with free cysteines (27 in SDS-insoluble and 96 in SDS-soluble), 330 RmCys peptides (110 in SDS-insoluble and 230 in SDS-soluble). Additionally, 72 non-modified proteins were differentially regulated (45 in SDS-insoluble and 30 in SDS-soluble), representing less than 2% of the total proteome – consistent with expectations.

Further analyzing these changes, most differentially regulated formerly SA N-glycopeptides were associated with cell adhesion (e.g., THBS1, MCAM and LAMA4), synaptic components (e.g., SV2B, LAMP5), and proteins involved in neurite outgrowth and synapse formation (e.g., SERPINE2, SEMA3C, GPRIN1).

Similarly, regulated phosphoproteins were associated with cytoskeletal dynamics, growth cones and synaptogenesis. These included DCX and GAP43 – both also altered in the synaptic structure phosphoproteome -, as well as NECTIN3, RTN4, CTTN, and MINK1.

Differentially regulated peptides with free cysteines mapped to proteins linked to growth cone biology, SV cycle and synapse-associated extracellular matrix (ECM), in addition to cytoskeleton and microtubule motor activity. Examples included MAP1B, NCAM1 – well-known for its role in neuron-neuron adhesion and neurite outgrowth-, VCAN, and NRP2-receptor for different class of semaphorins-, CLTA – major component of coated vesicles - and EXOC3-4 - exocyst complex components and SV docking. Proteins regulated in RmCys - representing for example disulfide bonds or S-nitrosylation - were similarly implicated in ECM (e.g., DST, MFAP2), cytoskeletal regulation, and synapse formation and growth cone dynamics, including ICAM5, TNC, and SLIT2. Interestingly, several Wnt pathway components - WNT8B, WNT7A, FZD7 and DKK1- were also differentially regulated by RmCys modifications.

In summary, a total of 526 proteins in the P1 fraction displayed SCZ-associated changes in PTMs. Overall, functional analysis revealed alterations of factors that support growth cone dynamics (including axonogenesis and neurite outgrowth) (Figure 4B), cytoskeletal organization and cell-cell interactions (including anchoring junctions, cell adhesion, and cadherin binding), and synapse structure and activity (e.g., PSD, SVs) (Supplementary Figure S4).

### Functional characterization of synaptic structures

Since neurotransmitter release from synapses are strongly regulated by phosphorylation-dependent mechanisms, we investigated the phosphorylation changes in proteins in SCZ-and control-derived synaptic structures in response to acute depolarization (15 sec), induced by high concentration of KCl. Synapse structure from SCZ and controls purified by differential centrifugation was treated with KCl or vehicle and the quantitative changes in phosphorylation were assessed using TiO_2_ chromatography and TMT-based quantitation. Subsequently, changes in phosphorylation were analyzed by bioinformatics tools, to identify shared and condition-specific phosphoprotein regulations (Figure 5A).

**Figure 5.**
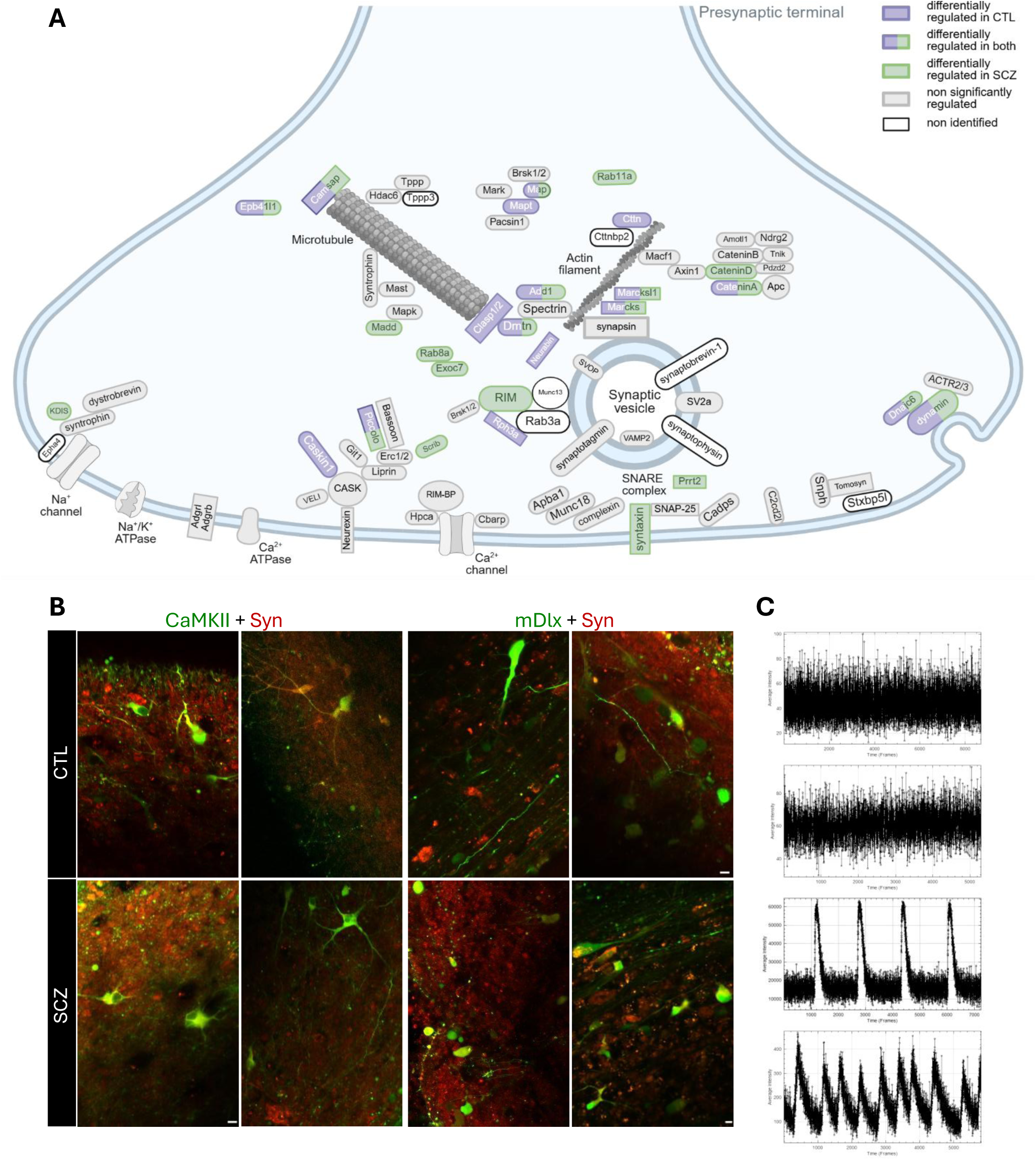
A.Schematic overview of differentially regulated phosphoproteins of SCZ and CTL synaptosomes in response to KCl stimulation **B.** Representative z-projections of calcium imaging fields of view (FOVs) from CTL-and SCZ-derived ALI-FOs transduced with calcium indicators targeting glutamatergic (CaMKII, green), GABAergic (mDlx, green) and general neuronal (Syn, red) population. Scale bar = 10 µm **C.** Representative calcium traces from individual regions of interest (ROI), showing spontaneous excitatory and inhibitory activity over time, and more complex firing patterns (bottom plots).

A total of 10864 phosphopeptides were quantified, of which 89 were significantly regulated in CTL and 157 in SCZ upon depolarization, corresponding to 77 and 107 proteins, respectively. The shared stimulation response is represented by 35 phosphoproteins, including key components of the AZ and SV cycle, such as PCLO, DNM1, CAMK2D, and MAP2. For proteins like MARCKSL1, CHGA, and ADD1 we observed different phosphosites differentially regulated between SCZ and control. Among the phospho-signaling event identified, DNM1 showed statistically significant dephosphorylation – monophosphorylated S774 and diphosphorylated S774 and S778 peptides – in both CTL-and SCZ-derived synaptic structures after depolarization (Supplementary Figure S2D). These changes, consistent with the role of DNM1 in depolarization-dependent endocytosis, confirm the functional integrity of enriched synaptic structures, as well as effectiveness of the acute KCl-induced depolarization.

CTL-specific regulated proteins were predominantly associated with cytoskeletal organization and dynamics, including CTTN, MAPT, CLASP2, and L1CAM. In contrast, SCZ-specific KCl response was primarily linked to SV cycle - encompassing both exocytosis and endocytosis mechanisms – as well as cytoskeletal dynamics. Key proteins involved in these pathways included STX1A/B, RAB8A, RIMS1, MADD, SCRIB, TRIOBP, and NEFM. Notably, consistent with previously mentioned observations, several proteins strongly associated with axon guidance, cytoskeletal rearrangement, and growth cone dynamics were also found regulated upon depolarization. Specifically, these included DPYSL2, ENAH, CRMP1, MYH10 in both SCZ and CTL response, DPYSL3, NFASC, CHL1, and ABI2 only in SCZ, and MYH9 and L1CAM uniquely in CTL.

### Calcium Imaging

Between days 170 and 190 the presence of spontaneous neuronal activity was assessed in both SCZ and CTL ALI-FOs using calcium imaging. To enable a more comprehensive functional characterization, two commercial viruses expressing GCAMP fluorescent calcium indicator - a genetically encoded calcium sensor that fluoresces green upon Ca^2+^ binding - were individually employed to selectively target glutamatergic and GABAergic neuronal populations (CaMKII and mDlx promoters, respectively). These were simultaneously transduced with rGECO indicator - that fluoresces red in response to Ca^2+^ binding - targeting the entire neuronal population (Syn promoter). These enabled the optical study of calcium-dependent processes, allowing the reconstruction of glutamatergic and GABAergic calcium dynamics with millisecond resolution. ALI-FOs from both conditions exhibited spontaneous excitatory and inhibitory activity. As expected, all glutamatergic and GABAergic neurons expressed also the Syn promoter; however, not all syn-positive neurons belonged to these subtypes – as indicated by the presence of red-only cells (Figure 5B). Additionally, more complex firing patterns – e.g., neuronal synchronicity-were observed in a subset of samples (Figure 5C).

## Discussion

In this study, we investigated synaptic molecular function and its influence on SCZ pathogenesis using human ALI-FOs generated from iPSCs reprogrammed from PBMCs derived from individuals with SCZ and healthy controls. Synaptic structures were enriched using our optimized protocol for ALI_FOs ^39^, and their proteomic and phosphoproteomic profiles, as well as their capacity of depolarization-induced phospho-signaling, were characterized by Mass Spectrometry. In parallel, we conducted a broader analysis of the PTMome of the P1 fraction, which includes the surrounding cellular environment and additional synaptic structures not released by the homogenization in the hypotonic buffer. Furthermore, we assessed spontaneous calcium activity of glutamatergic and GABAergic neuronal populations in the ALIFOs after day 170, using cell-type-specific viral transduction of calcium indicators.

### Impaired synaptogenesis and neurite outgrowth

SCZ pathogenesis is widely accepted to originate during prenatal brain development, as proposed by the neurodevelopmental hypothesis^53–55^. However, the molecular mechanisms underlying this process remain poorly understood. Growing evidence supports the involvement of early alterations in neuronal development^56–58^, synaptogenesis^30,54^ and synaptic activity^15,59,60^ in SCZ. Building on these previous findings and considering the limited availability of human fetal brain tissue, we generated ALI-FOs derived from iPSCs from individuals with SCZ and CTL to model human early brain development and synaptogenesis.

SCZ-and CTL-derived ALI-FOs exhibited comparable morphological architecture throughout the entire development. Extended culturing up to 170 days ensured the maturation of ALI-FOs, including the formation of functional synapses and neural circuits^19^. On day 170 the presence of key cellular components - neurons, synapses, mitochondria, and astrocytes - was confirmed through IHC labeling, while maturation stage and synaptic structure integrity were further validated by TEM images of enriched synaptic structures. Notably, TEM images of both conditions revealed the presence of mature synaptic structures, characterized by numerous SVs of similar size, mitochondria, and specialized areas (e.g., AZ, PSD) identifiable as electron-dense regions. In addition, SCZ-derived organoids exhibited growth cones, distinguishable by the presence of filopodia, and an emptier and granular cytoplasm almost completely lacking SVs and specialized areas. These features in SCZ may reflect ongoing or delayed synaptogenic activity.

To investigate synapse-specific molecular content, synaptic structures were enriched and further fractionated into SDS-soluble and SDS-insoluble samples, followed by MS analysis. A previous study has shown that a small concentration of SDS detergent in combination with ultracentrifugation could be used to specifically separate the AZ - detergent insoluble - in isolated synaptosomes^50,61^. This approach enabled the detection of nearly 1000 additional proteins compared to other fractions (P1 and S1), while specifically enriching synaptic proteins, both from pre-and post-synaptic terminals. Consistent with prior observations in SCZ^9,11^, the overall overlap in protein identities between SCZ and CTL-derived proteomes exceeded 98% across all analyzed fractions (P1, S1, SDS-soluble and SDS-insoluble P5000), and only a small percentage of proteins and phosphoproteins was differentially regulated between the conditions.

Our data showed that the majority of the dysregulated proteins and phosphoproteins were functionally grouped into three strongly interconnected categories: (1) synaptogenesis and synapse signaling; (2) cytoskeleton and cell junctions; (3) growth cone dynamics and neurite outgrowth. Specifically, we identified alterations in key molecular factors implicated in growth cone guidance signaling and establishment of neuron-neuron interactions – e.g., depletion of SEMA3E and CNTN5, and regulation of DCX and DCC at the phosphorylation level. In addition, cytoskeletal components and regulators - such as MARCKS, NEFL, STMN1, and MYO10 – were also affected. These proteins are known to be essential for motility, neurite outgrowth and axon pathfinding^4,62,63^. Notably, alterations in NEFL, STMN1, and MYO10 have previously been reported in several SCZ postmortem brain tissues across multiple brain regions, including corpus callosum, anterior and posterior cingulate cortex, and dorsolateral prefrontal cortex^4,64–69^.

Likewise, supporting the hypothesis of altered synaptogenesis, we also observed significant phospho-regulation of neuromodulin, a major marker of growth cones^70–73^ - but also a persistently expressed Ca^2+^-sensitive protein in presynaptic site of mature synapses^74,75^. Depletion of neuromodulin, as well as reduced levels of its mRNA, has been widely associated with SCZ in both postmortem brain – across different brain regions, including the frontal cortex, hippocampus, and primary visual cortex - and *in vitro* studies, with decreased levels already during neurodevelopment^11,67,76–78^. Similarly, levels of neuropilin 1 (NRP1) - another axonal growth cone marker^70^ – were elevated in the SCZ-derived P5000 fraction, further suggesting the potential impairment of synapse formation in SCZ-derived organoids.

Lastly, we detected a significant depletion of Wnt signaling components in SCZ-derived synaptosomes, encompassing WLS, WNT5A, WNT5B, WNT3, WNT8B, SFRP2, and LGR4. These changes indicate potential disruptions of Wnt biogenesis and trafficking, as well as impairments of both Wnt canonical and non-canonical pathways at SCZ synapses. WNTs are secreted glycoproteins, signaling through at least three divergent downstream pathways – canonical pathway, planar cell polarity (PCP) pathway, and Wnt/calcium pathway - upon binding to Frizzled receptors, and occasionally to Ryk (receptor tyrosine kinase) receptors, and requiring Dishevelled (Dvl) cytoplasmic proteins^79,80^. WNTs can act both locally - autocrine or adjacent cells – and at distance – e.g., by generating signaling gradients^81^. Wnt pathway functions are diverse and partly unknown, made complex by region-specificity and cell compartment-specificity^82^. It is well known that Wnt signaling is broadly implicated in several mechanisms of brain development, including synaptogenesis, axon guidance, neuroplasticity, and neurogenesis^80–84^. For instance, during axon guidance and synapse formation, WNTs regulate axon remodeling and branching - including their microtubule rearrangements and stability - and axon pathfinding, promoting the outgrowth through both repulsive or attractive cues and signaling gradients to axon navigation coordination^80,81^. Additionally, it has been observed that growth cone size and branching are also modulated by WNTs, as well as synapse assembly^80^. During synapse formation, WNTs are required for clustering of synaptic components within growth cones prior to the establishment of pre-and post-synapse contact, as well as for the subsequent crosstalk between pre-and post-synaptic sites to ensure coordinated synapse assembly^81,83^. Wnt pathway is highly controlled during early brain development, and its loss in function has been observed to lead to defects in localization, morphology and integrity of synaptic components, as well as disassemble of synapse and synaptic plasticity deficits^80,83,85^.

Interestingly, Wnt signaling impairment has been associated with several neuropsychiatric and neurodegenerative disorders including SCZ, bipolar disorder, and ASD^79,86–91^. Given the complexity of Wnt signaling and its implication in several mechanisms, its specific role in SCZ pathology and pathogenesis is still poorly understood^92^. However, SCZ postmortem brain tissues, blood samples, and animal model studies have reported diverse Wnt pathway components and downstream proteins to be affected^88^. Specifically, changes in protein levels, mRNA and enzymatic activity of glycogen synthase kinase 3 beta (GSK3) have been observed across different brain regions^93^, along with altered expression of β-and γ-catenin, WNT1, and FZD7 receptor, and GSK3 phosphorylation state^79,86,92,93^. Furthermore, WNT5a non-canonical signaling pathway has been observed altered in SCZ^94^, as well as various Wnt antagonists, such as DKK1, DKK3, and DKK4^79,92^. Notably, in support of the involvement of Wnt pathway in SCZ, it has been observed that administration of both typical (e.g., haloperidol) and atypical (e.g., risperidone, clozapine) antipsychotics used in SCZ affects several Wnt signaling components, leading to altered levels of GSK3, β-catenin, Dvl, WNT5a, and Axin in different brain regions^87,95,96^.

### PTMome analysis support synaptic alterations

Building upon the alterations identified in the synaptic structure fraction, we extended our investigation to the broader PTMome of the P1 fraction, which includes the surrounding cellular environment of synapses and additional synaptic structures not released from the cell body by the homogenization in hypotonic buffer. This focus on PTMs, rather than protein abundance, was driven by the limited proteomic changes previously observed in SCZ^9,11^, and by the subsequent hypothesis that differences in regulation of protein activity, rather than expression, may play a more central role in the overall function of synapses. Herein this context, PTMs refer to the analysis of protein phosphorylation, peptides with free cysteines, peptides with reversibly modified cysteines, and sialylated N-linked glycopeptides, all protein features capable of modulate protein function, activity and interaction in cellular signaling pathways^51,52^.

Proteins carrying significantly regulated PTMs in SCZ were again primarily associated with growth cone dynamics and neurite outgrowth – e.g., SERPINE2, SEMA3C, NRP2 GPRIN1, SIRPA, CTTN-, cytoskeletal organization and cell-cell interactions – e.g., NCAM1, MAP1B, ICAM5, TNC, MCAM, LAMA4-, and synaptic structure and activity – e.g., SV2B, CLTA, RAB8A, EXOC7, EXOC3. These findings support the alterations observed in the synaptic structure fractions, highlighting the potential role of PTMs in SCZ-related synaptic dysfunction.

Notably, key growth cone and neurodevelopmental markers - GAP43 and DCX - were again differentially regulated in the P1 fraction, in agreement with our previous observations in the P5000 synaptic structure fraction. Similarly, FABP7, a protein previously reported as highly abundant in growth cones^71,72^, showed a significant regulation in P1. Likewise, additional proteins known to be highly enriched in growth cones, even if also present in mature synaptosomes^70,73^ - such as HSP90AB1, MAP1B, RTN4, LRP1, DYNC1H1, and NEFM – were regulated in P1. Among these, MAP1B has been identified as the primary structural component of the growth cone cytoskeleton^71,73^, whose activity is strictly regulated by phosphorylation^73,97^. Similarly, the motor protein dynein heavy chain 1 (DYNC1H1), essential for vesicular transport, has been reported as one of the most abundant proteins in growth cones^70^.

Finally, dysregulation of the Wnt signaling pathway was also detected within the P1 fraction, involving a different set of components - WNT8B, WNT7A, FZD7 and DKK1-suggesting a broader alteration of the Wnt pathway in SCZ.

### KCl-induced phospho-signaling

Synaptosomes are known to retain full viability after enrichment, preserving ATP generation, metabolic and enzymatic activity, membrane potential, ion homeostasis, and intact neurotransmitter machinery^98–100^. Thus, synaptosomes are suitable for functional studies of molecular mechanisms underlying synaptic transmission ^98,101,102^. Likewise, growth cones retain their immature but functional uptake and release machinery – potentially involved in modulating neuronal development, but not in synaptic signal transmission - as well as the capability of depolarization response to a certain extent^103,104^.

Here, we evaluated the response of SCZ-and CTL-derived synaptic structures to acute KCl-induced depolarization by characterizing changes in phosphorylation signaling. It is well established that synaptic signal transmission is tightly controlled by phosphatase and kinase activity (e.g., Calcium/calmodulin-dependent protein kinase II (CaMKII) and calcineurin), which regulates multiple phosphoproteins involved in SV exo-and endo-cytosis^100,105,106^.

As previously described, depolarization triggers the influx of Ca^2+^ through voltage-gated ion channels, inducing phosphorylation and dephosphorylation of multiple presynaptic proteins that regulate the neurotransmitter release apparatus^100,105^. Depolarization-dependent signaling in both SCZ-and CTL-derived synaptic structures involved phosphorylation changes in key components of the AZ and SV cycle, including proteins such as PCLO, DNM1, DPYSL2, DNAJC6, MARCKS, and MAP2. Interestingly, among these phospho-signaling events, DNM1 – known to be a key regulator of SV endocytosis^107,108^– revealed statistically significant dephosphorylation in both the identified monophosphorylated (S774) and diphosphorylated (S774 and S778) peptides. It has been established that upon acute depolarization – such as 15 seconds 76 mM KCl-induced stimulation - activity-dependent bulk endocytosis is activated within synaptic structures^109,110^, occurring in close temporal and spatial coupling with exocytosis^111^. A hallmark of activity-dependent bulk endocytosis is the rapid dephosphorylation of DNM1 at S774 and S778 by calcineurin, triggering its translocation - particularly its fast-responding short isoform (DNM1b) - to the plasma membrane^108,112^. SV retrieval is then initiated by self-assembly of DNM1 into collar-like structure around the endocytic invagination neck^112^ and by the recruitment of other dephosphins (e.g., amphiphysin I/II, synaptojanin, epsin, AP180, syndapin I) promoted by their coordinated dephosphorylation^108–110^. Thus, these data confirm the presence of active synaptic structures and successful depolarization in both CTL-and SCZ-derived enriched fractions.

Furthermore, many of these regulated phosphoproteins (22 out of 35) were reported in previous studies^100,105^. These overlapped phospho-signals included PCLO, CRMP1, DNAJC6, NCAM, DMTN, and DPYSL2. However, the extent and nature of the shared response may be influenced by differences in experimental conditions, including the developmental stage of the starting material - ALI-FOs contained less mature and developing synaptic structures - as well as a lower density of synaptic structures in our enriched fractions. Additionally, given the rapid dynamics of depolarization-dependent phosphorylation, differences in stimulation duration may also have affected the observed changes. Finally, as these experiments have not been performed in human synapses before, species-specific differences are expected when comparing responses between mouse and human synaptic structures.

Interestingly, CTL-specific phospho-response was mostly enriched in cytoskeleton-related proteins, such as CTTN, MAPT, and CLASP2. In contrast, SCZ-specific depolarization response prominently involved proteins associated with SV cycle mechanisms and synaptic function, including well-characterized proteins such as syntaxin-1A and B (STX1A/B), RIMS1, SCRIB, and RAB8A. Functional annotation of these regulated proteins revealed enrichment in pathways related to synaptic signaling, vesicle trafficking, and cytoskeletal rearrangement - in line with previous functional study of synaptosomes^105^. Finally, consistent with the presence of growth cones in our synaptic structure fractions, and with their known responsiveness to chemical stimulation^103,113,114^, the phospho-signaling profile also included several proteins highly expressed in growth cones and implicated in axon guidance process^71,72^ - e.g., GAP43, NCAM1, DPYSL2, MAP1B, CRMP1.

### Calcium activity

To further investigate the level of synaptic maturation and the development of self-organized neuronal networks in CTL and SCZ ALI-FOs, we characterized spontaneous excitatory and inhibitory neural activity using calcium imaging. Both SCZ and CTL ALI-FOs displayed glutamatergic and GABAergic activity, indicative of mature neuronal morphology and functional synaptic transmission, in line with previous findings in NOs^19,115^. In addition, more complex firing patterns - especially synchronized activity - were recorded in a subset of organoids from both conditions. These patterns suggest the emergence of mature neuronal networks, in which GABAergic neurons play a crucial role^116^. Indeed, inhibitory neurons are essential for modulating network neuroactivity, and their dysfunction has been shown to disrupt neural synchrony^117^.

In our study the presence of GABAergic neurons in both CTL and SCZ ALI-FOs at day 170 was also supported by proteomic analysis, which detected GABAergic markers, including calretinin (CALB2) and somatostatin (SST), as well as GABA receptor subunits (e.g., GABRA1-3, GABRB2, GABRD) and GABA transporters (e.g., SLC6A1, SLC6A11, SLC6A13).

SCZ has been consistently associated with disruptions in neural circuitry, including impaired neural synchrony – attributed to dysfunctional connectivity and altered GABAergic signalling^36,117,118^. Thus, further investigations exploring inhibitory neuron function and network activity differences in SCZ are needed. The use of ALI-FOs may provide a fundamental opportunity to uncover dysfunctional connectivity occurring during early brain development and to distinguish alterations from later stages of SCZ pathogenesis.

Together, our findings provide novel insight into early synaptic dysfunction in SCZ, highlighting molecular and functional impairments in growth cone dynamics, cytoskeletal organization, and synaptic signaling using patient iPSC derived ALI-FOs. These alterations align with converging evidence proposing synaptic dysfunction as a central feature of SCZ pathology, and support the hypothesis that SCZ originates, at least in part, during early neurodevelopment.

While our results contribute to deepening our knowledge of SCZ mechanisms during early brain development, future studies are needed. In particular, larger donor cohorts, additional functional characterization, and further validations will be essential.

## Acknowledgements

We would like to thank Prof. Kim Rewitz and his group for kindly providing access to his cell culture laboratory. This research was financially supported by the Lundbeck foundation collaborative grant DEVELOPNOID (R336-2020-1113) and the Villum Centre for Bioanalytical Sciences at SDU. This project was further supported by a grant from the Danish Agency of Higher Education and Science to establish the PLATO research infrastructure: Danish National Mass Spectrometry Platform for Proteomics and Biomolecular Imaging (grant no. 5229-00012B, www.sdu.dk/PLATO).

## Supplementary Figures

**Figure S1.**
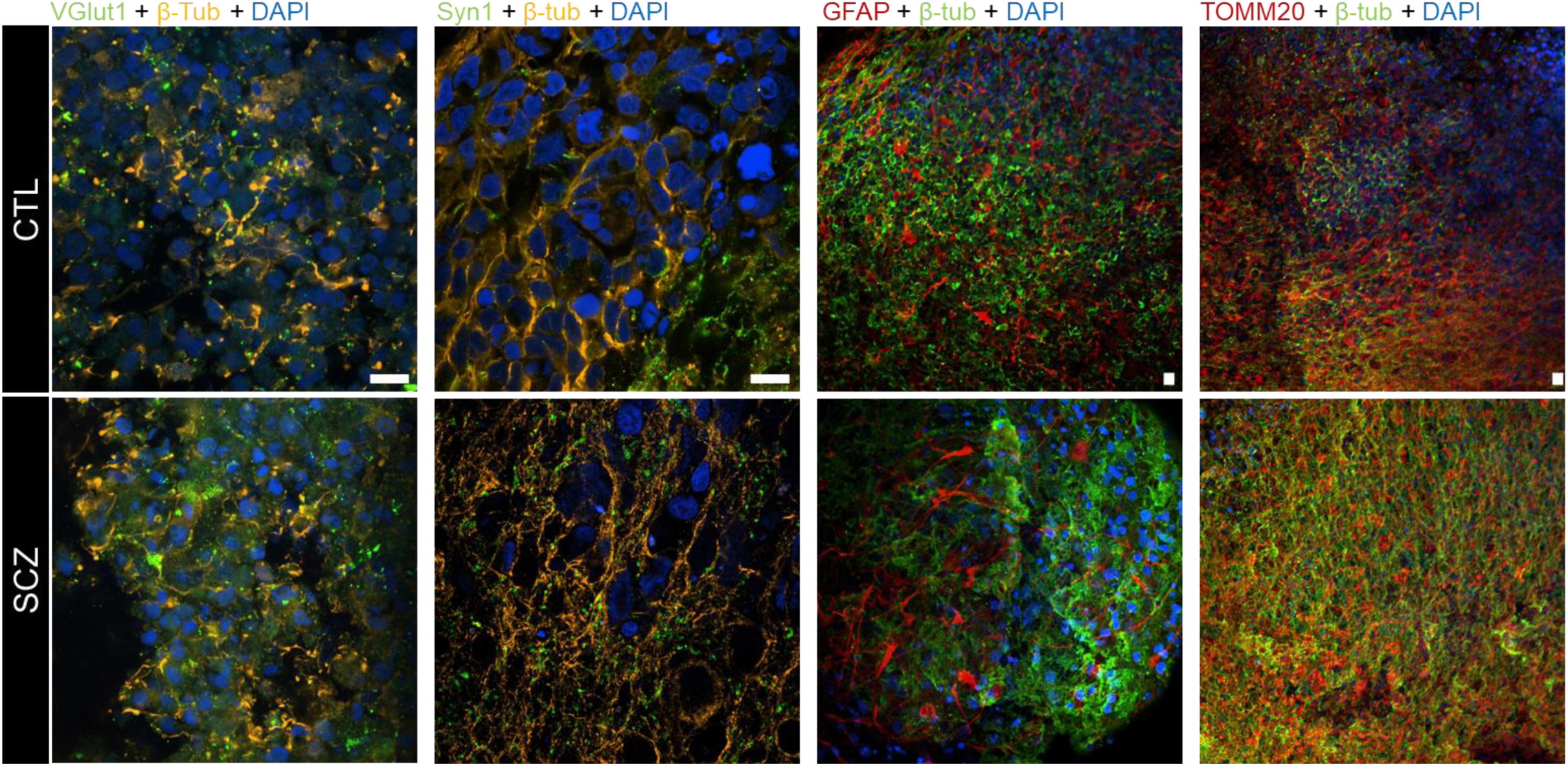
Immunohistochemical labelling of CTL and SCZ ALI-FOs at day 170 for vesicular glutamate transporter 1 (VGlut1, green), beta tubulin (β-Tub, orange or green), synapsin 1 (Syn1, green), glial fibrillary acidic protein (GFAP, red), mitochondrial import receptor subunit TOM20 homolog (TOMM20, red), with DAPI (dark blue). Scale bar = 10 µm

**Figure S2.**
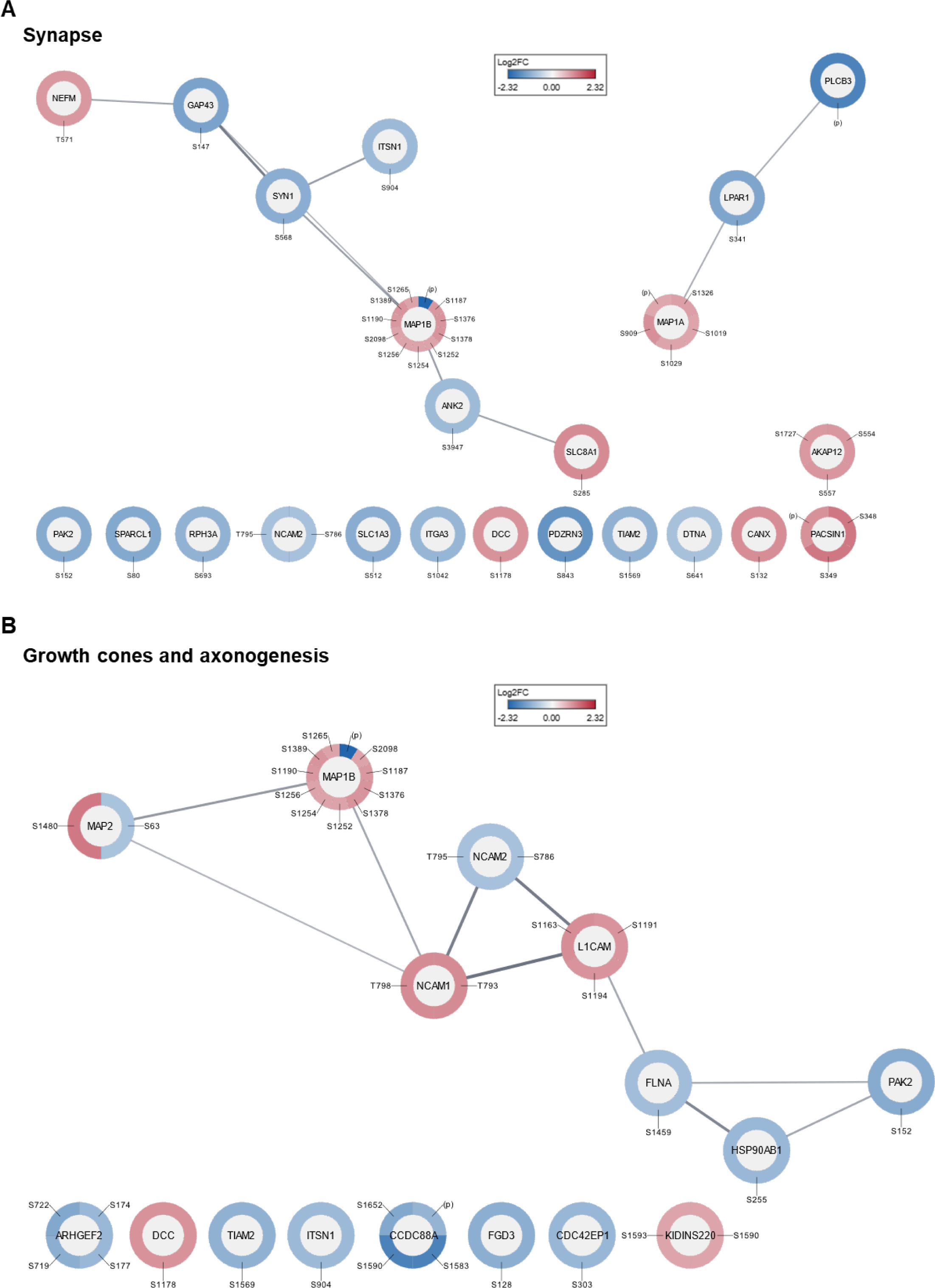
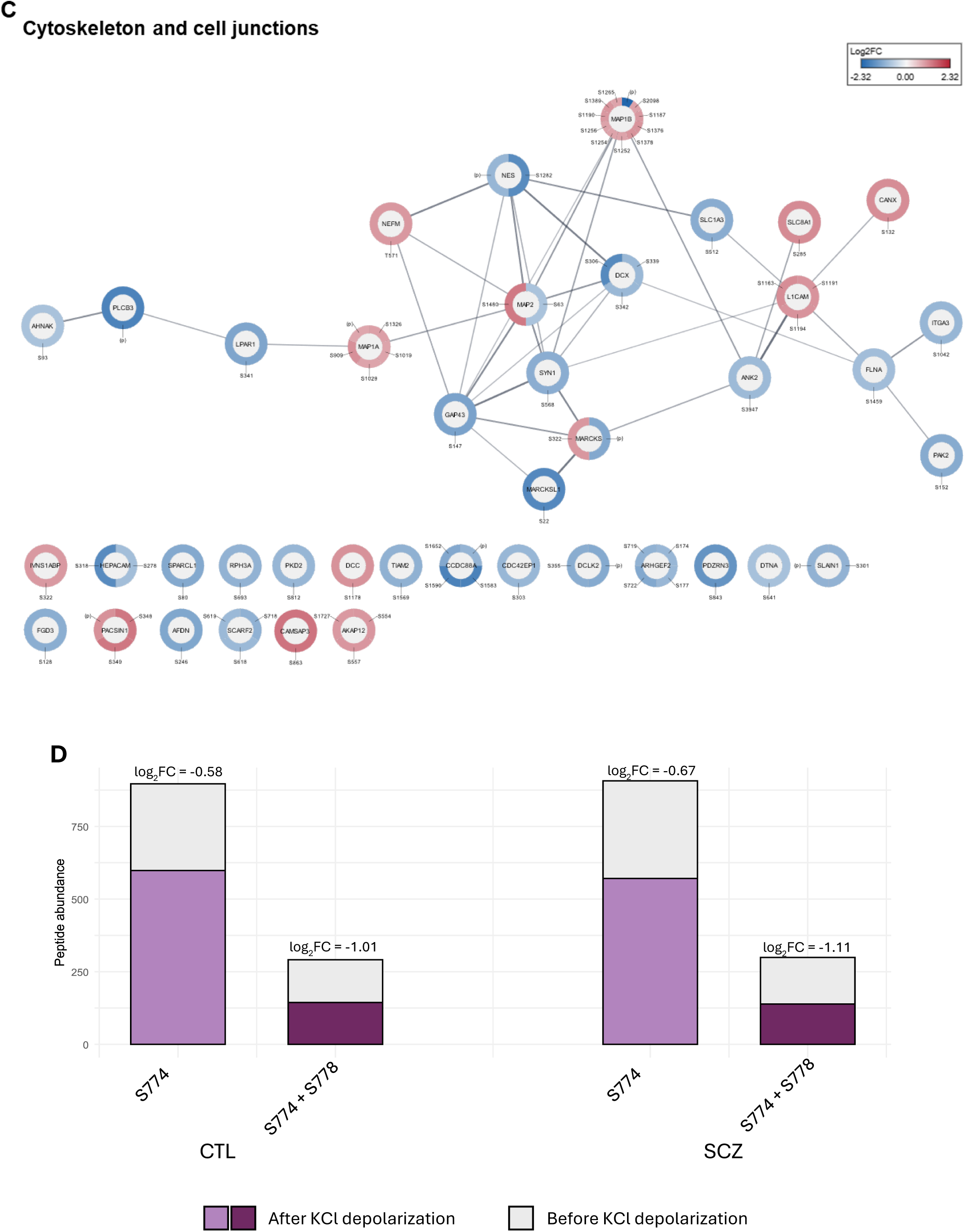
String networks of differentially regulated phosphoproteins in SCZ synaptosomes compared to CTL, grouped per GO terms – **A.** Synapses, **B.** Growth cones and axonogenesis, and **C.** Cytoskeleton and cell junctions. **D.** Bar plot illustrating the statistically significant (adj p-value < 0.05) dephosphorylation of DNM1 monophosphorylated (S774) and diphosphorylated (S774 + S778) peptides in response to KCl-induced depolarization in synaptic structures derived from CTL and SCZ ALI-FOs.

**Figure S3.**
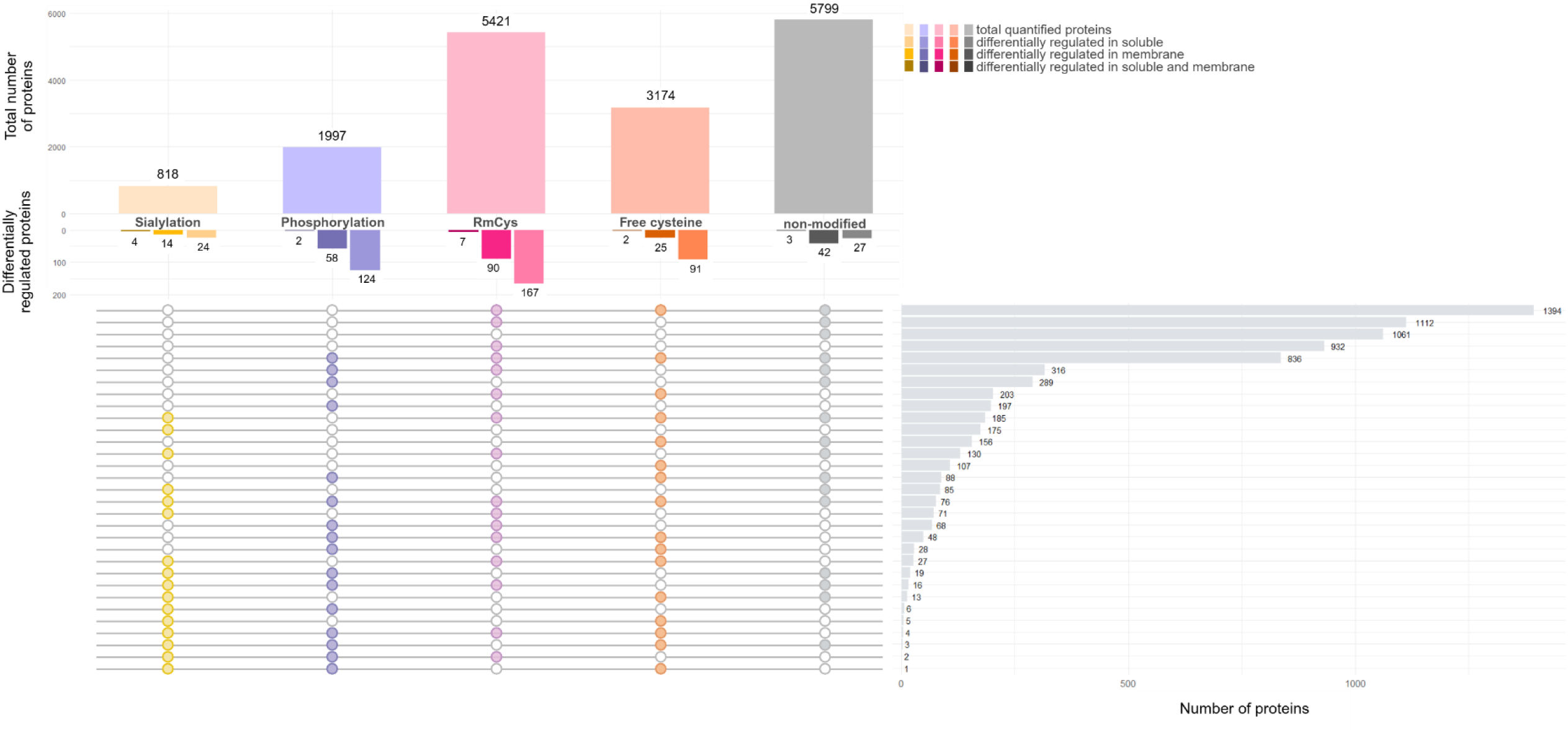
Overview of post-translationally modified peptides detected in the P1 fraction. The top bar plot shows the number of quantified proteins for each PTM investigated, as well as non-modified peptides. The bottom bar plot displays the number of differentially regulated proteins in SCZ compared to CTL. The lower part of the graph illustrates the overlap between proteins quantified in PTMomics and proteomics.

**Figure S4.**
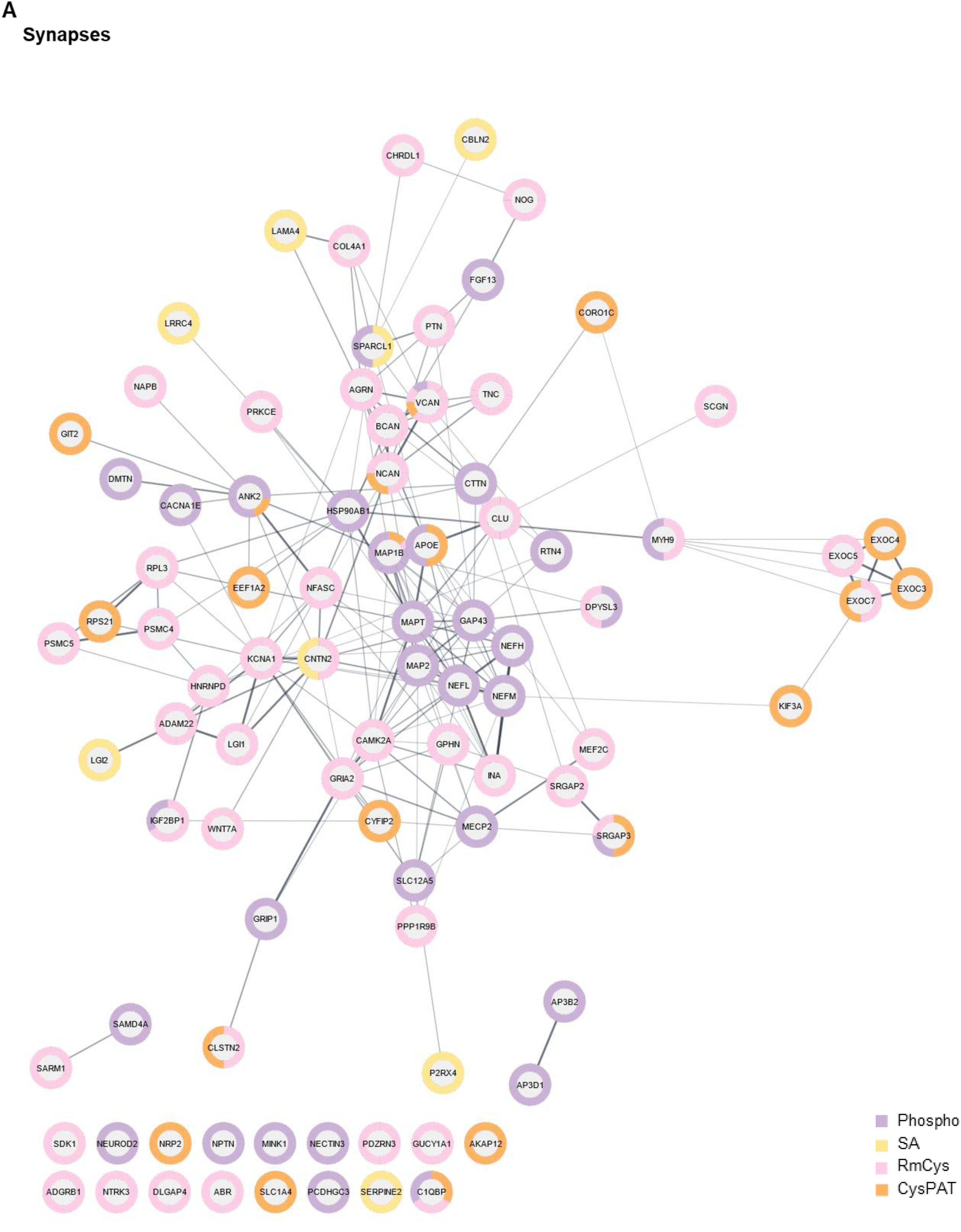
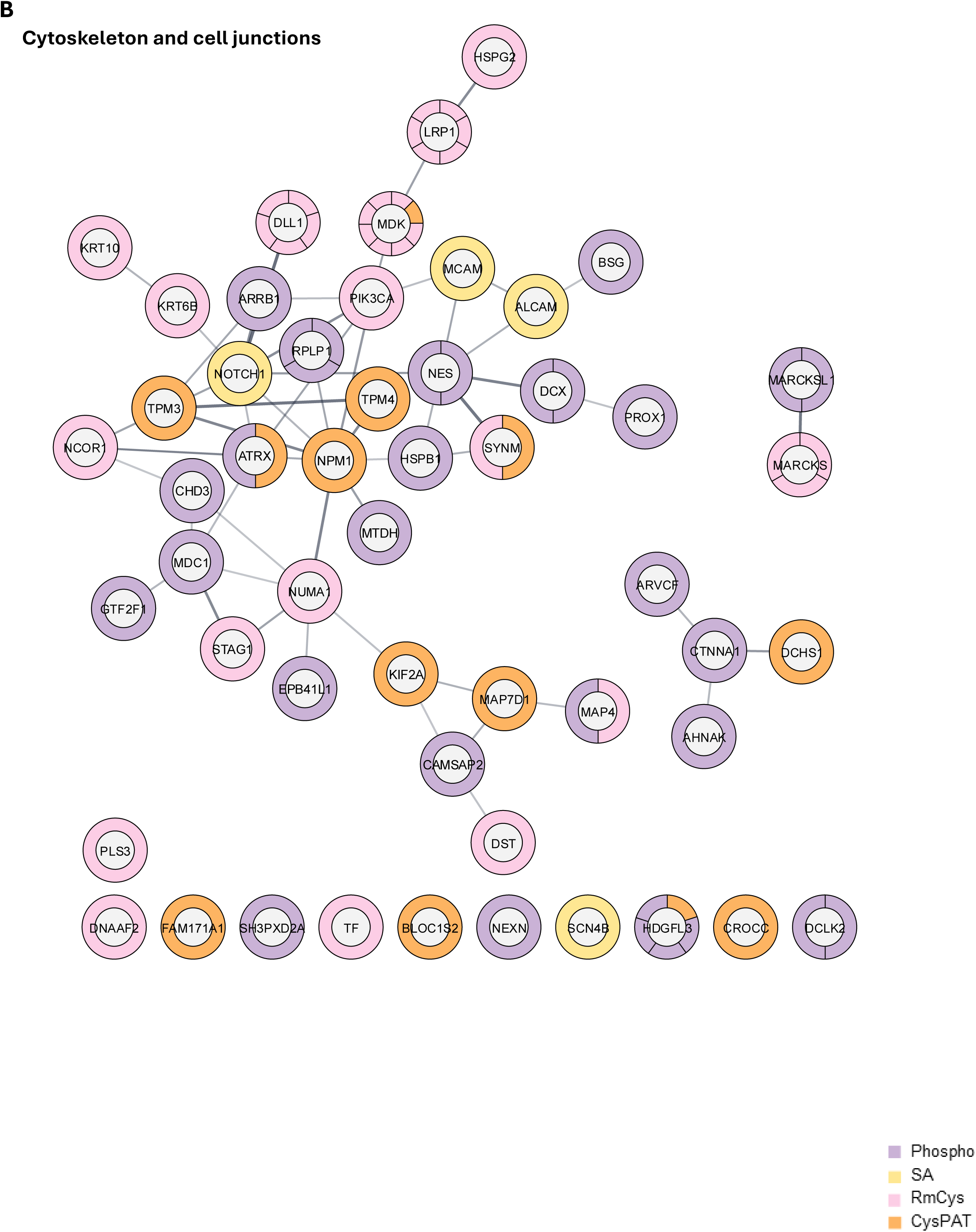
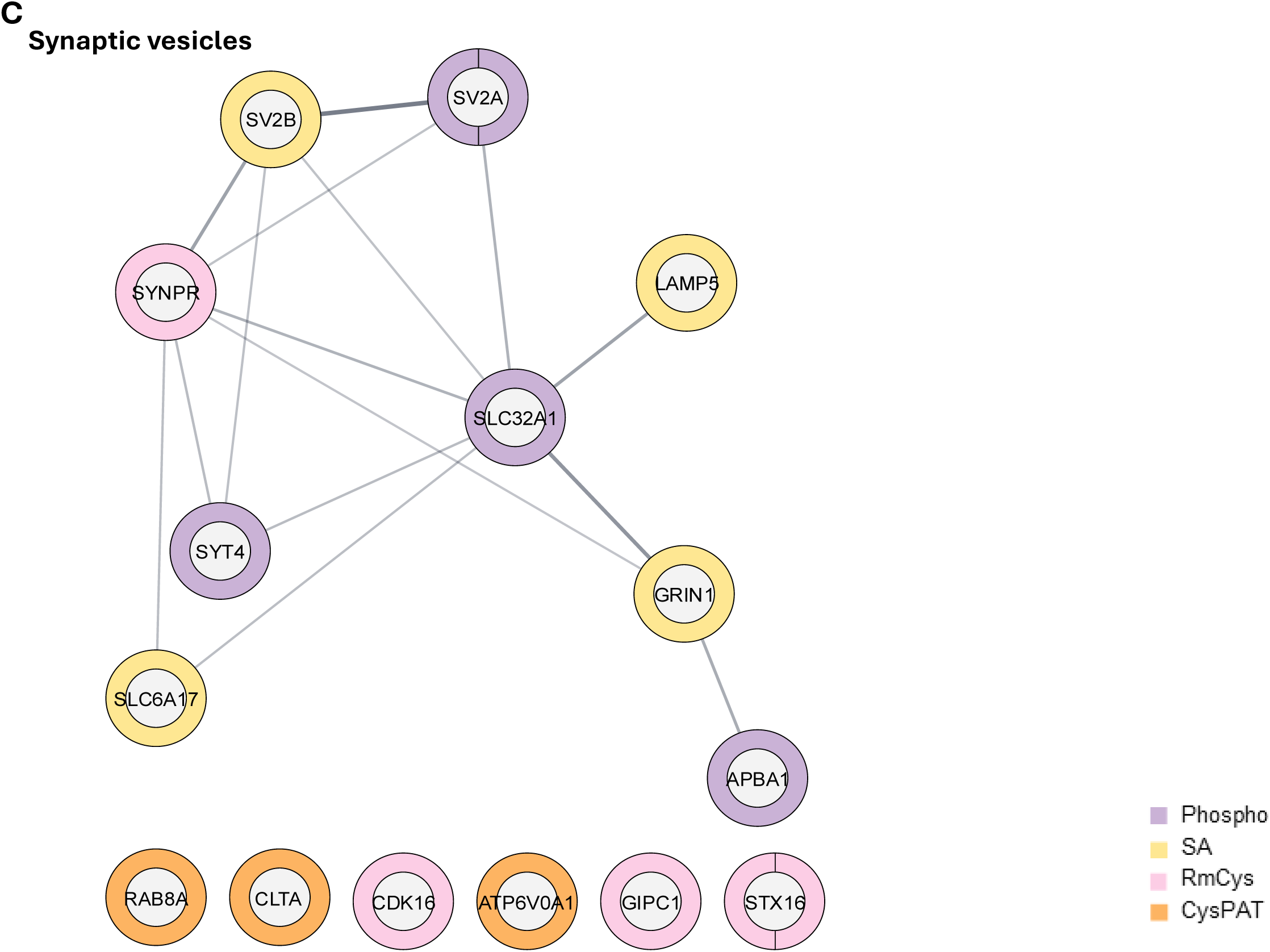
String networks of proteins with differentially regulated post<u>-</u>translationally modified peptides detected in SCZ-derived P1 compared to CTL. Networks were grouped per GO terms – **A.** Synapses, **B.** Cytoskeleton and cell junctions, **C.** Synaptic vesicles. Modifications include sialylation (yellow), phosphorylation (violet), reversed modified cysteine (RmCys, pink), and free cysteines (orange)

Synaptosome_KClstimulation_Phosphopeptides.xlsx

**Table S1** Phosphoproteomics on CTL and SCZ synaptosomes after KCl stimulation or control buffer incubation.

A. All identified phosphopeptides

B. Differentially regulated phosphopeptides in CTL synaptosomes in response to KCl depolarization

C. Differentially regulated phosphopeptides in SCZ synaptosomes in response to KCl depolarization

## References

1 Li, X. et al. The global burden of schizophrenia and the impact of urbanization during 1990-2019: An analysis of the global burden of disease study 2019. Environ Res 232, 116305 (2023). 10.1016/j.envres.2023.116305

2 Kahn, R. S. et al. Schizophrenia. Nature Reviews Disease Primers 1, 15067 (2015). 10.1038/nrdp.2015.67

3 Steiner, J., Guest, P. C. & Martins-de-Souza, D. in Proteomic Methods in Neuropsychiatric Research (ed Paul C. Guest) 3-19 (Springer International Publishing, 2017).

4 Nascimento, J. M. & Martins-de-Souza, D. The proteome of schizophrenia. npj Schizophrenia 1, 14003 (2015). 10.1038/npjschz.2014.3

5 Howes, O. D. & Onwordi, E. C. The synaptic hypothesis of schizophrenia version III: a master mechanism. Molecular Psychiatry 28, 1843–1856 (2023). 10.1038/s41380-023-02043-w

6 Chung, Y. & Cannon, T. D. Brain Imaging During the Transition from Psychosis Prodrome to Schizophrenia. The Journal of Nervous and Mental Disease 203 (2015).

7 Martin Bauer, N. P.-R. S. K. M. W. Is Dopamine Neurotransmission Altered in Prodromal Schizophrenia? A Review of the Evidence. Current Pharmaceutical Design 18, 1568–1579 (2012). 10.2174/138161212799958611

8 Yilmaz, M. et al. Overexpression of schizophrenia susceptibility factor human complement C4A promotes excessive synaptic loss and behavioral changes in mice. Nature Neuroscience 24, 214–224 (2021). 10.1038/s41593-020-00763-8

9 Notaras, M. et al. Schizophrenia is defined by cell-specific neuropathology and multiple neurodevelopmental mechanisms in patient-derived cerebral organoids. Molecular Psychiatry 27, 1416–1434 (2022). 10.1038/s41380-021-01316-6

10 Nagy, C. et al. Effects of Postmortem Interval on Biomolecule Integrity in the Brain. Journal of Neuropathology & Experimental Neurology 74, 459–469 (2015). 10.1097/NEN.0000000000000190

11 Notaras, M., Lodhi, A., Fang, H., Greening, D. & Colak, D. The proteomic architecture of schizophrenia iPSC-derived cerebral organoids reveals alterations in GWAS and neuronal development factors. Translational Psychiatry 11, 541 (2021). 10.1038/s41398-021-01664-5

12 Aryal, S., et al. Deep proteomics identifies shared molecular pathway alterations in synapses of patients with schizophrenia and bipolar disorder and mouse model. Cell Reports 42 (2023). 10.1016/j.celrep.2023.112497

13 Stachowiak, E. K. et al. Cerebral organoids reveal early cortical maldevelopment in schizophrenia—computational anatomy and genomics, role of FGFR1. Translational Psychiatry 7, 6 (2017). 10.1038/s41398-017-0054-x

14 Dubonyte, U., Asenjo-Martinez, A., Werge, T., Lage, K. & Kirkeby, A. Current advancements of modelling schizophrenia using patient-derived induced pluripotent stem cells. Acta Neuropathologica Communications 10, 183 (2022). 10.1186/s40478-022-01460-2

15 Trubetskoy, V. et al. Mapping genomic loci implicates genes and synaptic biology in schizophrenia. Nature 604, 502–508 (2022). 10.1038/s41586-022-04434-5

16 Nucifora, F. C., Woznica, E., Lee, B. J., Cascella, N. & Sawa, A. Treatment resistant schizophrenia: Clinical, biological, and therapeutic perspectives. Neurobiology of Disease 131, 104257 (2019). 10.1016/j.nbd.2018.08.016

17 Birtele, M., Lancaster, M. & Quadrato, G. Modelling human brain development and disease with organoids. Nat Rev Mol Cell Biol 26, 389–412 (2025). 10.1038/s41580-024-00804-1

18 Qian, X., Song, H. & Ming, G.-l. Brain organoids: advances, applications and challenges. Development 146, dev166074 (2019). 10.1242/dev.166074

19 Giandomenico, S. L. et al. Cerebral organoids at the air–liquid interface generate diverse nerve tracts with functional output. Nature Neuroscience 22, 669–679 (2019). 10.1038/s41593-019-0350-2

20 Qian, X. et al. Brain-Region-Specific Organoids Using Mini-bioreactors for Modeling ZIKV Exposure. Cell 165, 1238–1254 (2016). 10.1016/j.cell.2016.04.032

21 Jo, J. et al. Midbrain-like Organoids from Human Pluripotent Stem Cells Contain Functional Dopaminergic and Neuromelanin-Producing Neurons. Cell Stem Cell 19, 248–257 (2016). 10.1016/j.stem.2016.07.005

22 Jgamadze, D. et al. Protocol for human brain organoid transplantation into a rat visual cortex to model neural repair. STAR Protocols 4, 102470 (2023). 10.1016/j.xpro.2023.102470

23 Di Lullo, E. & Kriegstein, A. R. The use of brain organoids to investigate neural development and disease. Nature Reviews Neuroscience 18, 573–584 (2017). 10.1038/nrn.2017.107

24 Vanova, T., et al. Cerebral organoids derived from patients with Alzheimer’s disease with PSEN1/2 mutations have defective tissue patterning and altered development. Cell Reports 42 (2023). 10.1016/j.celrep.2023.113310

25 Smits, L. M. et al. Modeling Parkinson’s disease in midbrain-like organoids. npj Parkinson’s Disease 5, 5 (2019). 10.1038/s41531-019-0078-4

26 Raja, W. K. et al. Self-Organizing 3D Human Neural Tissue Derived from Induced Pluripotent Stem Cells Recapitulate Alzheimer’s Disease Phenotypes. PLOS ONE 11, e0161969 (2016). 10.1371/journal.pone.0161969

27 Antón-Bolaños, N. et al. Brain Chimeroids reveal individual susceptibility to neurotoxic triggers. Nature 631, 142–149 (2024). 10.1038/s41586-024-07578-8

28 Krenn, V. et al. Organoid modeling of Zika and herpes simplex virus 1 infections reveals virus-specific responses leading to microcephaly. Cell Stem Cell 28, 1362–1379.e1367 (2021). 10.1016/j.stem.2021.03.004

29 Yang, A. C. & Tsai, S.-J. New Targets for Schizophrenia Treatment beyond the Dopamine Hypothesis. International Journal of Molecular Sciences 18 (2017).

30 Habela, C. W., Song, H. & Ming, G.-l. Modeling synaptogenesis in schizophrenia and autism using human iPSC derived neurons. Molecular and Cellular Neuroscience 73, 52–62 (2016). 10.1016/j.mcn.2015.12.002

31 MacDonald, M. L. et al. Synaptic Proteome Alterations in the Primary Auditory Cortex of Individuals With Schizophrenia. JAMA Psychiatry 77, 86–95 (2020). 10.1001/jamapsychiatry.2019.2974

32 Moyer, C. E., Shelton, M. A. & Sweet, R. A. Dendritic spine alterations in schizophrenia. Neuroscience Letters 601, 46–53 (2015). 10.1016/j.neulet.2014.11.042

33 Brennand, K. J. et al. Modelling schizophrenia using human induced pluripotent stem cells. Nature 473, 221–225 (2011). 10.1038/nature09915

34 Ziermans, T. B. et al. Progressive structural brain changes during development of psychosis. Schizophr Bull 38, 519–530 (2012). 10.1093/schbul/sbq113

35 Berdenis van Berlekom, A., et al. Synapse Pathology in Schizophrenia: A Meta-analysis of Postsynaptic Elements in Postmortem Brain Studies. Schizophrenia bulletin 46 (2019). 10.1093/schbul/sbz060

36 Spencer, K. M. et al. Abnormal Neural Synchrony in Schizophrenia. The Journal of Neuroscience 23, 7407 (2003). 10.1523/JNEUROSCI.23-19-07407.2003

37 Avram, M., Brandl, F., Bäuml, J. & Sorg, C. Cortico-thalamic hypo-and hyperconnectivity extend consistently to basal ganglia in schizophrenia. Neuropsychopharmacology 43, 2239–2248 (2018). 10.1038/s41386-018-0059-z

38 Comer, A. L. et al. Increased expression of schizophrenia-associated gene C4 leads to hypoconnectivity of prefrontal cortex and reduced social interaction. PLOS Biology 18, e3000604 (2020). 10.1371/journal.pbio.3000604

39 Ohlenschlaeger, M. S. et al. Modeling Synaptic Maturation From Growth Cone to Synapse in Human Organoids. J Neurochem 170, e70458 (2026). 10.1111/jnc.70458

40 Konala, V. B. R., Nandakumar, S., Surendran, H. & Pal, R. in Induced Pluripotent Stem Cells and Human Disease: Methods and Protocols (ed Kursad Turksen) 137–151 (Springer US, 2022).

41 Ashrafian, S. et al. Quantitative Phosphoproteomics and Acetylomics of Safranal Anticancer Effects in Triple-Negative Breast Cancer Cells. Journal of Proteome Research 21, 2566–2585 (2022). 10.1021/acs.jproteome.2c00168

42 Huang, H. et al. Simultaneous Enrichment of Cysteine-containing Peptides and Phosphopeptides Using a Cysteine-specific Phosphonate Adaptable Tag (CysPAT) in Combination with titanium dioxide (TiO_2_) Chromatography *. Molecular & Cellular Proteomics 15, 3282–3296 (2016). 10.1074/mcp.M115.054551

43 Lucrezia Criscuolo, S. B. E., Arkadiusz Nawrocki, Lene A. Jakobsen, Pia Jensen, Peter T. Jensen, Honggang Huang, Jesper F. Havelund, Nils J. Færgeman, Helle Bogetofte, Martin R. Larsen. MOSAIC-PTM: A Sequential Enrichment of Multiple Protein Post-Translational Modifications with Isobaric Tag-Based Multiplexing for Comprehensive PTM Profiling. Submitted to Mol. Cell. Proteomics (2026).

44 Spivak, M., Weston, J., Bottou, L., Käll, L. & Noble, W. S. Improvements to the percolator algorithm for Peptide identification from shotgun proteomics data sets. J Proteome Res 8, 3737–3745 (2009). 10.1021/pr801109k

45 Wu, T. et al. clusterProfiler 4.0: A universal enrichment tool for interpreting omics data. The Innovation 2 (2021). 10.1016/j.xinn.2021.100141

46 Koopmans, F. et al. SynGO: An Evidence-Based, Expert-Curated Knowledge Base for the Synapse. Neuron 103, 217–234.e214 (2019). 10.1016/j.neuron.2019.05.002

47 Wickham, H. ggplot2: Elegant Graphics for Data Analysis. 2 edn, (Springer Cham, 2016).

48 Legeay, M., Doncheva, N., Morris, J. & Jensen, L. Visualize omics data on networks with Omics Visualizer, a Cytoscape App [version 2; peer review: 2 approved]. F1000Research 9 (2020). 10.12688/f1000research.22280.2

49 Shannon, P. et al. Cytoscape: a software environment for integrated models of biomolecular interaction networks. Genome Res 13, 2498–2504 (2003). 10.1101/gr.1239303

50 Boll, I., Jensen, P., Schwammle, V. & Larsen, M. R. Depolarization-dependent Induction of Site-specific Changes in Sialylation on N-linked Glycoproteins in Rat Nerve Terminals. Mol Cell Proteomics 19, 1418–1435 (2020). 10.1074/mcp.RA119.001896

51 Leutert, M., Entwisle, S. W. & Villén, J. Decoding Post-Translational Modification Crosstalk With Proteomics. Mol Cell Proteomics 20, 100129 (2021). 10.1016/j.mcpro.2021.100129

52 Venne, A. S., Kollipara, L. & Zahedi, R. P. The next level of complexity: Crosstalk of posttranslational modifications. PROTEOMICS 14, 513–524 (2014). 10.1002/pmic.201300344

53 Birnbaum, R. & Weinberger, D. R. The Genesis of Schizophrenia: An Origin Story. Am J Psychiatry 181, 482–492 (2024). 10.1176/appi.ajp.20240305

54 Schmitt, A., Falkai, P. & Papiol, S. Neurodevelopmental disturbances in schizophrenia: evidence from genetic and environmental factors. J Neural Transm (Vienna) 130, 195–205 (2023). 10.1007/s00702-022-02567-5

55 McGrath, J. J., P., F. F., J., B. T. H., Alan, M.-S. & and Eyles, D. W. The neurodevelopmental hypothesis of schizophrenia: a review of recent developments. Annals of Medicine 35, 86–93 (2003). 10.1080/07853890310010005

56 Reif, A. et al. Neural stem cell proliferation is decreased in schizophrenia, but not in depression. Mol Psychiatry 11, 514–522 (2006). 10.1038/sj.mp.4001791

57 Na, K. S., Jung, H. Y. & Kim, Y. K. The role of pro-inflammatory cytokines in the neuroinflammation and neurogenesis of schizophrenia. Prog Neuropsychopharmacol Biol Psychiatry 48, 277–286 (2014). 10.1016/j.pnpbp.2012.10.022

58 Iannitelli, A., Quartini, A., Tirassa, P. & Bersani, G. Schizophrenia and neurogenesis: A stem cell approach. Neurosci Biobehav Rev 80, 414–442 (2017). 10.1016/j.neubiorev.2017.06.010

59 Malhotra, D. et al. High frequencies of de novo CNVs in bipolar disorder and schizophrenia. Neuron 72, 951–963 (2011). 10.1016/j.neuron.2011.11.007

60 Glessner, J. T. et al. Strong synaptic transmission impact by copy number variations in schizophrenia. Proc Natl Acad Sci U S A 107, 10584–10589 (2010). 10.1073/pnas.1000274107

61 Matthies, I. et al. N-Glycosylation in isolated rat nerve terminals. Mol Omics 17, 517–532 (2021). 10.1039/d0mo00044b

62 Dent, E. W., Gupton, S. L. & Gertler, F. B. The growth cone cytoskeleton in axon outgrowth and guidance. Cold Spring Harb Perspect Biol 3 (2011). 10.1101/cshperspect.a001800

63 Gordon-Weeks, P. R. & Fournier, A. E. Neuronal cytoskeleton in synaptic plasticity and regeneration. J Neurochem 129, 206–212 (2014). 10.1111/jnc.12502

64 Sivagnanasundaram, S., Crossett, B., Dedova, I., Cordwell, S. & Matsumoto, I. Abnormal pathways in the genu of the corpus callosum in schizophrenia pathogenesis: a proteome study. PROTEOMICS – Clinical Applications 1, 1291–1305 (2007). 10.1002/prca.200700230

65 Martins-de-Souza, D. et al. Alterations in oligodendrocyte proteins, calcium homeostasis and new potential markers in schizophrenia anterior temporal lobe are revealed by shotgun proteome analysis. Journal of Neural Transmission 116, 275–289 (2009). 10.1007/s00702-008-0156-y

66 Reis-de-Oliveira, G. et al. Digging deeper in the proteome of different regions from schizophrenia brains. Journal of Proteomics 223, 103814 (2020). 10.1016/j.jprot.2020.103814

67 Föcking, M. et al. Proteomic and genomic evidence implicates the postsynaptic density in schizophrenia. Molecular Psychiatry 20, 424–432 (2015). 10.1038/mp.2014.63

68 Saia-Cereda, V. M. et al. Proteomics of the corpus callosum unravel pivotal players in the dysfunction of cell signaling, structure, and myelination in schizophrenia brains. European Archives of Psychiatry and Clinical Neuroscience 265, 601–612 (2015). 10.1007/s00406-015-0621-1

69 Martins-de-Souza, D. et al. Sex-specific proteome differences in the anterior cingulate cortex of schizophrenia. Journal of Psychiatric Research 44, 989–991 (2010). 10.1016/j.jpsychires.2010.03.003

70 Estrada-Bernal, A. et al. Functional Complexity of the Axonal Growth Cone: A Proteomic Analysis. PLOS ONE 7, e31858 (2012). 10.1371/journal.pone.0031858

71 Nozumi, M. et al. Identification of functional marker proteins in the mammalian growth cone. Proceedings of the National Academy of Sciences 106, 17211–17216 (2009). 10.1073/pnas.0904092106

72 Chauhan, M. Z. et al. Multi-Omic Analyses of Growth Cones at Different Developmental Stages Provides Insight into Pathways in Adult Neuroregeneration. iScience 23 (2020). 10.1016/j.isci.2020.100836

73 Igarashi, M. Proteomic identification of the molecular basis of mammalian CNS growth cones. Neuroscience Research 88, 1–15 (2014). 10.1016/j.neures.2014.07.005

74 Gamby, C., Waage, M. C., Allen, R. G. & Baizer, L. Analysis of the Role of Calmodulin Binding and Sequestration in Neuromodulin (GAP-43) Function*. Journal of Biological Chemistry 271, 26698–26705 (1996). 10.1074/jbc.271.43.26698

75 Liu, Y. C. & Storm, D. R. Dephosphorylation of Neuromodulin by Calcineurin. Journal of Biological Chemistry 264, 12800–12804 (1989). 10.1016/S0021-9258(18)51557-X

76 Chambers, J. S., Thomas, D., Saland, L., Neve, R. L. & Perrone-Bizzozero, N. I. Growth-associated protein 43 (GAP-43) and synaptophysin alterations in the dentate gyrus of patients with schizophrenia. Progress in Neuro-Psychopharmacology and Biological Psychiatry 29, 283–290 (2005). 10.1016/j.pnpbp.2004.11.013

77 Sower, A. C., Bird, E. D. & Perrone-Bizzozero, N. I. Increased levels of GAP-43 protein in schizophrenic brain tissues demonstrated by a novel immunodetection method. Molecular and Chemical Neuropathology 24, 1–11 (1995). 10.1007/BF03160108

78 Weickert, C. S. et al. Reduced GAP-43 mRNA in Dorsolateral Prefrontal Cortex of Patients with Schizophrenia. Cerebral Cortex 11, 136–147 (2001). 10.1093/cercor/11.2.136

79 Mulligan, K. A. & Cheyette, B. N. R. Neurodevelopmental Perspectives on Wnt Signaling in Psychiatry. Molecular Neuropsychiatry 2, 219–246 (2017). 10.1159/000453266

80 Rosso, S. B. & Inestrosa, N. C. WNT signaling in neuronal maturation and synaptogenesis. Front Cell Neurosci 7, 103 (2013). 10.3389/fncel.2013.00103

81 Ciani, L. & Salinas, P. C. WNTS in the vertebrate nervous system: from patterning to neuronal connectivity. Nature Reviews Neuroscience 6, 351–362 (2005). 10.1038/nrn1665

82 Valvezan, A. J. & Klein, P. S. GSK-3 and Wnt Signaling in Neurogenesis and Bipolar Disorder. Frontiers in Molecular Neuroscience Volume 5–2012 (2012). 10.3389/fnmol.2012.00001

83 Ahmad-Annuar, A. et al. Signaling across the synapse: a role for Wnt and Dishevelled in presynaptic assembly and neurotransmitter release. Journal of Cell Biology 174, 127–139 (2006). 10.1083/jcb.200511054

84 Varela-Nallar, L., Alfaro, I. E., Serrano, F. G., Parodi, J. & Inestrosa, N. C. Wingless-type family member 5A (Wnt-5a) stimulates synaptic differentiation and function of glutamatergic synapses. Proceedings of the National Academy of Sciences 107, 21164–21169 (2010). 10.1073/pnas.1010011107

85 Marzo, A. et al. Reversal of Synapse Degeneration by Restoring Wnt Signaling in the Adult Hippocampus. Curr Biol 26, 2551–2561 (2016). 10.1016/j.cub.2016.07.024

86 Cotter, D., et al. Abnormalities of Wnt signalling in schizophrenia – evidence for neurodevelopmental abnormality. NeuroReport 9 (1998).

87 Singh, K. K. An emerging role for Wnt and GSK3 signaling pathways in schizophrenia. Clinical Genetics 83, 511–517 (2013). 10.1111/cge.12111

88 Hoseth, E. Z. et al. Exploring the Wnt signaling pathway in schizophrenia and bipolar disorder. Translational Psychiatry 8, 55 (2018). 10.1038/s41398-018-0102-1

89 Srikanth, P. et al. Genomic *DISC1* Disruption in hiPSCs Alters Wnt Signaling and Neural Cell Fate. Cell Reports 12, 1414–1429 (2015). 10.1016/j.celrep.2015.07.061

90 Belinson, H. et al. Prenatal β-catenin/Brn2/Tbr2 transcriptional cascade regulates adult social and stereotypic behaviors. Molecular Psychiatry 21, 1417–1433 (2016). 10.1038/mp.2015.207

91 Srikanth, P. et al. Shared effects of DISC1 disruption and elevated WNT signaling in human cerebral organoids. Translational Psychiatry 8, 77 (2018). 10.1038/s41398-018-0122-x

92 Karabicici, M., Azbazdar, Y., Iscan, E. & Ozhan, G. Misregulation of Wnt Signaling Pathways at the Plasma Membrane in Brain and Metabolic Diseases. Membranes (Basel) 11 (2021). 10.3390/membranes11110844

93 Sutton, L. P., Honardoust, D., Mouyal, J., Rajakumar, N. & Rushlow, W. J. Activation of the canonical Wnt pathway by the antipsychotics haloperidol and clozapine involves dishevelled-3. Journal of Neurochemistry 102, 153–169 (2007). 10.1111/j.1471-4159.2007.04527.x

94 Evgrafov, O. V. et al. Gene Expression in Patient-Derived Neural Progenitors Implicates WNT5A Signaling in the Etiology of Schizophrenia. Biol Psychiatry 88, 236–247 (2020). 10.1016/j.biopsych.2020.01.005

95 Beaulieu, J.-M., Gainetdinov, R. R. & Caron, M. G. Akt/GSK3 Signaling in the Action of Psychotropic Drugs. Annual Review of Pharmacology and Toxicology 49, 327–347 (2009). 10.1146/annurev.pharmtox.011008.145634

96 Proitsi, P. et al. Positional Pathway Screen of wnt Signaling Genes in Schizophrenia: Association with DKK4. Biological Psychiatry 63, 13–16 (2008). 10.1016/j.biopsych.2007.03.014

97 Riederer, B. M. Microtubule-associated protein 1B, a growth-associated and phosphorylated scaffold protein. Brain Research Bulletin 71, 541–558 (2007). 10.1016/j.brainresbull.2006.11.012

98 Daniel, J. A., Malladi, C. S., Kettle, E., McCluskey, A. & Robinson, P. J. Analysis of synaptic vesicle endocytosis in synaptosomes by high-content screening. Nature Protocols 7, 1439–1455 (2012). 10.1038/nprot.2012.070

99 Jhou, J.-F. & Tai, H.-C. The Study of Postmortem Human Synaptosomes for Understanding Alzheimer’s Disease and Other Neurological Disorders: A Review. Neurology and Therapy 6, 57–68 (2017). 10.1007/s40120-017-0070-z

100 Engholm-Keller, K. et al. The temporal profile of activity-dependent presynaptic phospho-signalling reveals long-lasting patterns of poststimulus regulation. PLoS Biol 17, e3000170 (2019). 10.1371/journal.pbio.3000170

101 Ahmad, F., Jing, Y., Lladó, A. & Liu, P. Chemical Stimulation of Rodent and Human Cortical Synaptosomes: Implications in Neurodegeneration. Cells 10 (2021).

102 Bradford, H. F. Metabolic response of synaptosomes to electrical stimulation: Release of amino acids. Brain Research 19, 239–247 (1970). 10.1016/0006-8993(70)90437-3

103 Lockerbie, R. O. Biochemical pharmacology of isolated neuronal growth cones: implications for synaptogenesis. Brain Research Reviews 15, 145–165 (1990). 10.1016/0165-0173(90)90016-H

104 Lockerbie, R. O. et al. Cyclic AMP reduces adhesion of isolated neuronal growth cones from developing rat forebrain to an astrocytic cell line from embryonic mouse striatum. Neuroscience 28, 443–454 (1989). 10.1016/0306-4522(89)90191-7

105 Silbern, I. et al. Protein Phosphorylation in Depolarized Synaptosomes: Dissecting Primary Effects of Calcium from Synaptic Vesicle Cycling. Mol Cell Proteomics 20, 100061 (2021). 10.1016/j.mcpro.2021.100061

106 Cousin, M. A., Tan, T. C. & Robinson, P. J. Protein phosphorylation is required for endocytosis in nerve terminals: potential role for the dephosphins dynamin I and synaptojanin, but not AP180 or amphiphysin. J Neurochem 76, 105–116 (2001). 10.1046/j.1471-4159.2001.00049.x

107 Liu, J.-P., Sim, A. T. R. & Robinson, P. J. Calcineurin Inhibition of Dynamin I GTPase Activity Coupled to Nerve Terminal Depolarization. Science 265, 970–973 (1994). 10.1126/science.8052858

108 Anggono, V. et al. Syndapin I is the phosphorylation-regulated dynamin I partner in synaptic vesicle endocytosis. Nat Neurosci 9, 752–760 (2006). 10.1038/nn1695

109 Clayton, E. L., Evans, G. J. & Cousin, M. A. Bulk synaptic vesicle endocytosis is rapidly triggered during strong stimulation. J Neurosci 28, 6627–6632 (2008). 10.1523/jneurosci.1445-08.2008

110 Clayton, E. L. et al. The phospho-dependent dynamin-syndapin interaction triggers activity-dependent bulk endocytosis of synaptic vesicles. J Neurosci 29, 7706–7717 (2009). 10.1523/jneurosci.1976-09.2009

111 Cousin, M. A. & Robinson, P. J. Ca(2+) influx inhibits dynamin and arrests synaptic vesicle endocytosis at the active zone. J Neurosci 20, 949–957 (2000). 10.1523/jneurosci.20-03-00949.2000

112 Smillie, K. J. & Cousin, M. A. Dynamin I phosphorylation and the control of synaptic vesicle endocytosis. Biochem Soc Symp, 87–97 (2005). 10.1042/bss0720087

113 Saito, S., Komiya, Y. & Igarashi, M. Muscarinic acetylcholine receptors are expressed and enriched in growth cone membranes isolated from fetal and neonatal rat forebrain: Pharmacological demonstration and characterization. Neuroscience 45, 735–745 (1991). 10.1016/0306-4522(91)90285-V

114 Yamada, K. M., Spooner, B. S. & Wessells, N. K. ULTRASTRUCTURE AND FUNCTION OF GROWTH CONES AND AXONS OF CULTURED NERVE CELLS. Journal of Cell Biology 49, 614–635 (1971). 10.1083/jcb.49.3.614

115 Reumann, D. et al. In vitro modeling of the human dopaminergic system using spatially arranged ventral midbrain–striatum–cortex assembloids. Nature Methods 20, 2034–2047 (2023). 10.1038/s41592-023-02080-x

116 Sakaguchi, H. et al. Self-Organized Synchronous Calcium Transients in a Cultured Human Neural Network Derived from Cerebral Organoids. Stem Cell Reports 13, 458–473 (2019). 10.1016/j.stemcr.2019.05.029

117 Zhang, W. et al. Human forebrain organoid-based multi-omics analyses of PCCB as a schizophrenia associated gene linked to GABAergic pathways. Nature Communications 14, 5176 (2023). 10.1038/s41467-023-40861-2

118 Heesen, S. H. & Köhr, G. GABAergic interneuron diversity and organization are crucial for the generation of human-specific functional neural networks in cerebral organoids. Front Cell Neurosci 18, 1389335 (2024). 10.3389/fncel.2024.1389335

